# The mechanosensitive TRPV2 calcium channel controls human melanoma invasiveness and metastatic potential

**DOI:** 10.1101/2021.10.22.465391

**Authors:** Kenji F. Shoji, Elsa Bayet, Dahiana Le Devedec, Aude Mallavialle, Séverine Marionneau-Lambot, Sabrina Leverrier-Penna, Florian Rambow, Raul Perret, Aurélie Joussaume, Roselyne Viel, Alain Fautrel, Amir Khammari, Bruno Constantin, Sophie Tartare-Deckert, Aubin Penna

## Abstract

Discovery of therapeutic targets against metastasis is of primary importance since being the main cause of cancer-related death. Melanoma is a highly aggressive cancer endowed with a unique capacity of rapidly metastasizing. Deregulation of calcium homeostasis has been involved in numerous cellular metastatic behaviors, although the molecular determinants supporting these processes often remain unclear. Here, we evidenced a prominent expression of the plasma membrane TRPV2 calcium channel as a distinctive feature of melanoma tumors, directly related to melanoma metastatic progression and dissemination. *In vitro* as well as *in vivo*, TRPV2 activity was sufficient to confer both migratory and invasive phenotypes to non-invasive melanoma cells, while conversely upon TRPV2 silencing, highly metastatic melanoma cells failed to retain their malignant behaviors. We established a model whereupon activation of the mechanosensitive TRPV2 channel, localized in highly dynamic nascent adhesion clusters, directly regulates calpain-dependent cleavage of the adhesive protein talin together with F-actin network. By operating at the crossroad of the tumor microenvironment and the intracellular machinery, mechanosensitive TRPV2 channel controls melanoma cells aggressiveness. Finally in human melanoma tumor samples, TRPV2 overexpression represents a molecular marker of advanced malignancy and bad prognosis, highlighting a new therapeutic option for migrastatics in the treatment of metastatic melanoma.

**Significance:** One essential feature of metastatic cells is enhanced motility and invasiveness. This study evidences TRPV2 channel control over metastatic melanoma invasiveness, highlights new migration regulatory mechanisms, and reveals this channel as a biomarker and migrastatic target for the treatment of advanced melanoma.

## INTRODUCTION

Cutaneous malignant melanoma (CMM) is a cancer arising from skin melanocytes (1). *In situ* tumors can be cured by surgical resection, but melanoma has a distinct tendency to spread into multiple organs. Metastatic melanoma is the deadliest form of skin cancer, with a rising incidence (2). Over the last decade, immuno- and targeted-therapies have shown increasing clinical benefit, but remain often insufficient to achieve durable responses (1, 3). Hence, understanding the molecular mechanisms enabling the acquisition of the CMM pro-metastatic phenotype is critical for defining early biomarkers and novel therapeutic targets.

Melanomagenesis consists of an initial growth phase, with melanocytes proliferating within the primary site, then transitioning into an invasive slow-cycling phase where tumor cells spread, and ultimately resume proliferation to promote the growth of metastasis. The dynamic behavior of CMM dissemination is sustained by an increased cell motility and invasiveness, both requiring precise communication between cells and their environment to breach basement membranes and colonize surrounding tissues. Accumulating evidence has demonstrated that altered calcium (Ca^2+^) signaling promotes tumor cell-specific phenotypic changes, supporting the metastatic spread (4).

The transient receptor potential (TRP) and ORAI Ca^2+^ channels families have been identified as key actors of physiological and pathological cell migration and invasion (5, 6). These ion channels are emerging as very attractive therapeutic targets in oncology due to both their ability to switch on or off specific phenotypic hallmarks of tumor cells, and their accessibility to pharmacological modulation (7). Still, little is known on specific channel subunit input in the particular context of CMM progression (8, 9).

The scope of this study was to identify among the numerous Ca^2+^-conducting channels expressed in melanoma cells any atypical profile, to determine an eventual association with tumor metastatic progression, and to elucidate the associated molecular mechanisms. Using 2D and 3D *in vitro* models, as well as *in vivo* models and human tissues, we have established the essential role of the mechanosensitive Ca^2+^ channel TRPV2 during the metastatic switch of melanoma cells, defining this channel as a promising biomarker and migrastatic target.

## MATERIALS & METHODS

Please refer to SI for detailed protocols and additional information on cell culture, antibodies and chemicals.

### Mice model

All animal experiments were approved by the French animal care “comité d’éthique” in concordance with French and European Union law (license #44565) and conformed to the relevant regulatory standards. Experimental metastasis studies were performed as previously described (10, 11). Mice were housed at UTE-IRS1 (Nantes-University) under the animal care license #C44-278. Anaesthetized mice were placed into a restraining device, 10^6^ melanoma cells engineered to express a luciferase reporter gene (LUC cells) were suspended in 100 µL of PBS and injected through a 30-gauge needle into the tail vein of nude mice (scid hairless NOD (SHrN™) for 501mel, or NMRI for 451Lu).

### Plasmid constructs

The eGFP coding sequence from the pEGFP-C2 vector (Clontech), with or without the coding sequence of wild-type human TRPV2 (12) added in 3’, were inserted into the pcDNA3.1(+)-Zeocin vector (Life Technologies). All constructs were verified by sequencing. Validated non-silencing control and TRPV2 targeting shRNAmir-pGIPZ lentiviral vectors were purchased from Dharmacon.

### Lentiviral production and transfection/transduction procedures

Lentiviral particles production and cell transduction were performed according to manufacturer instructions. Briefly, 293SZ cells were co-transfected with shRNAmir-pGIPZ plasmids and the lentiviral psPAX2 and pMD2.G packaging plasmids in antibiotic-free medium using Ca^2+^-phosphate–mediated transfection. Target cells were infected with freshly thawed lentiviral particles diluted in growth medium supplemented with polybrene (3 μg/mL). For transfection, cells were electroporated with 10 μg of DNA using the BTM-830square waves generator (BTX Instrument Division, Harvard Apparatus). Selection of stable clones was achieved with selective doses of either zeocin (25 μg/mL) or puromycin for 2-4d, at which time mock-cells were eradicated. For the transfected 501mel cell lines, cell sorting based on GFP fluorescence was performed on a FACSAria Fusion cytometer (Becton Dickinson).

### Real-time quantitative PCR

Total RNA was isolated using the Nucleospin RNA II kit (Macherey-Nagel) following manufacturer’s instructions. RNA concentrations were estimated using a NanoDrop analyzer ND1000 (ThermoFisher). 0.5 μg purified RNA was reverse transcribed in a volume of 20 µL using the High-capacity cDNA Reverse Transcription kit (Applied Biosystems) and random hexamers according to manufacturer’s instructions. qPCR was performed on 0.5 ng cDNA samples, in sealed 96-well microtiter plates using the SYBR Green^TM^ PCR Master Mix (Applied Biosystems) and gene specific primer pairs (see Supplemental Table 1) with the 7300SDS Real-Time PCR System (Applied Biosystems). The ΔΔCt method was used to calculate relative expression values, which were normalized to the housekeeping gene Tbp.

### Bioinformatics analyses on publicly-available data

Data on TRPV2 transcript expression patterns in the NCI-60 cell line set were generated by querying TRPV2 as input in CellMiner (http://discover.nci.nih.gov/cellminer/) as described in (13). The NCI-60 is a panel of 60 diverse human cancer cell lines used by the Developmental Therapeutics Program of the U.S. National Cancer Institute RRID:SCR_003057.

The cancer genome atlas (TCGA) was interrogated for TRPV2 RNA expression across different tumor types and melanoma subgroups defined by the Clark level. Survival analysis (Kaplan-Meier estimate) was performed by comparing overall survival of the 10% highest to the 10% lowest TRPV2 expressers. The statistical significance was assessed with a Mantel-Cox log-rank test.

### Biochemical techniques

Immunoblotting was performed as previously described (14). The biotinylation assays were carried out using cell-impermeant EZ-Link™ Sulfo-NHS-LC-Biotin. (refer to SI for detailed procedures)

### Mn^2+^ quenching assay

Basal Ca^2+^ permeability was measured in Fura-2 loaded adherent melanoma cells. Briefly, cells were seeded in black 96-well clear bottom microplates (10^5^ cells/well), loaded with 5 µM Fura2/AM for 40 min at 37°C and washed with Ca^2+^-free HBSS solution (132 mM NaCl, 5.4 mM KCl, 0.8 mM MgCl_2_, 10 mM HEPES and 5.6 mM D-Glucose, pH7.4). A baseline was established in Ca^2+^-free HBSS solution then a final concentration of 215 µM MnCl_2_ was added. Fluorescence emission at 510 nm was acquired every 3s following Fura-2 excitation at its isosbestic point, 359 nm, on a multimode plate reader (Enspire, Perckin-Elmer). Mn^2+^ entry was measured as the rate of decline (quenching) of Fura-2 fluorescence intensity.

### Proximity ligation assay

PLA experiments were performed using the Red Mouse/Rabbit Duolink Starter Kit (Sigma-Aldrich) according to the manufacturer’s instructions. Plating, fixation, permeabilization and blocking were done as described above. F-actin was detected using Acti-stain-488-Phalloidin. Fluorescence was analyzed using the confocal microscope IX81-based Olympus FV1000 with UPLSAPO-NA 1.35 60X oil objective and the IQ3 software (Andor). Maximum projection intensities (MIP) of images were created from z stacks with a step interval of 0.2 μm. Quantification of PLA fluorescent spots was carried out using the particle analyzer application of ImageJ, RRID:SCR_003070 software.

### In vitro migration/invasion assays

Migration and invasion experiments were performed using transwell migration assays. Briefly, 2×10^5^ 501mel or WM266.4 cells, or 4×10^5^ 451Lu cells suspended in serum-free media were added to the top chamber of transwell permeable supports (8 μm pores, Corning) coated or not with Matrigel (Corning). Chemo-attraction was induced by media supplemented with 10% FCS into the bottom chamber. After 12h, cells were fixed in 70% cold ethanol and stained with Crystal Violet. Cells that have reached the downside of the porous membrane were counted using a cell counter (ImageJ).

### 2D cell migration and cell tracking

Serum-starved WM266.4 cells expressing shRNA control or TRPV2 were seeded in chambers coated with fibronectin (Ibidi) filled with serum-free medium. Migration was induced with a 5% FCS gradient and carried out for 12h. Pictures were taken using an inverted Olympus IX71 microscope equipped with a Cool SnapHQ camera installed on a Delta Vision system. Images were analyzed using the SofWorX and ImageJ software.

### Three-dimensional spheroid growth

Melanoma spheroids were generated using the liquid overlay technique and implanted into a type I collagen matrix in growth medium as previously described (15). Tumor cell outgrowth was visualized by phase contrast microscopy and the cell growth radius was measured with ImageJ.

### In vivo experiments

Metastasis formation and relative amounts of tumor burden were assessed weekly using whole-body bioluminescent imaging (BLI) (ΦimageurTM; Biospace Lab). Mice were given 150 mg/kg body-weight of D-luciferin potassium salt (Interchim). Images were acquired 3-5 min after injection and collected in real time until plate saturation was reached. Photons count per second per steradian per square centimeter was recorded by a photon imager system. For BLI plots, photon flux was calculated by using a rectangular ROI encompassing the thorax using the software Photovision+ (version 1.3; Biospace Lab). This value was normalized to the value obtained immediately after injection, so that all mice had an arbitrary starting signal of 100. At necropsy, *ex-vivo* BLI measurement was performed within 15 min after D-luciferin injection. Immunostaining procedure of lung metastases is detailed in SI.

### Immunohistochemistry

The melanoma tissue microarray ME1004c (US Biomax Inc.) came with the clinical stage, gender, age, organ, TNM classification and HMB45 profiles. Staining was performed on a Discovery Automated IHC stainer (Roche) using a rabbit polyclonal anti-TRPV2 antibody (1:100, HPA044993, RRID:AB_10960889), or the same concentration of a control isotype, or the melanoma triple cocktail (HMB45+A103+T311, Ventana, Roche). Signal enhancement was performed using the Ventana ChromoMap Kit Slides (biotin free system). Sections were counterstained with hematoxylin and mounted with DPX. A detailed assessment was done by anatomopathologists. Individual tumor regions were analyzed by color deconvolution using the Fast Red, Fast Blue, DAB filter (ImageJ, Fig. S13D) based on (16). Specific TRPV2 stained regions were then subjected to densitometry analysis. Optical density was obtained using the formula: OD= Log(255/Mean).

### Statistical analysis

All data are displayed as means ± SEM for n=3 biological replicates and are representative of at least three independently repeated experiments. Statistical differences among cell lines or treatments were done by paired Student *t* test or by ANOVA test for multiple comparisons, as appropriate. Statistical analyses were performed in Prism 6.0 (GraphPad Prism, RRID:SCR_002798). Values with a P value <0.05 were considered statistically significant.

### Data availability

The data generated in this study are available within the article and its supplementary data files. Further information and requests for resources and reagents may be directed to and will be fulfilled by the corresponding author.

## RESULTS

### TRPV2 is the predominantly expressed calcium channel in metastatic melanoma

To identify Ca^2+^-conducting channel subunits supporting melanoma metastasis formation, we screened the cancer genome atlas (TCGA) of skin cutaneous melanoma (SKCM) tumors. Among most members of the major Ca^2+^-permeable channel families, where at least one subunit has previously been associated with accrued motility behavior, TRPV2 transcript stood out as the uppermost expressed (Figure 1A). In a set of metastatic CMM cell lines, TRPV2 expression was likewise exceeding the other non-voltage gated Ca^2+^-channels levels (Figures 1B-C and data not shown). By querying transcriptomic data from the NCI-60 panel and the Broad-Novartis cancer cell line encyclopedia (CCLE), we evidenced that TRPV2 expression was distinctively exacerbated in melanoma cell lines, compared to all other cancer-derived cell lines (Figures 1D and S1A). At the protein level, the preferential distribution of TRPV2 within melanoma cells was clearly confirmed among a large panel of cancer cell lines originating from different tissues (Figure 1E). Importantly, the utmost expression of TRPV2 transcripts was also revealed in SKCM tumors compared to 36 other cancer types (Figure S1B).

**Figure 1.**
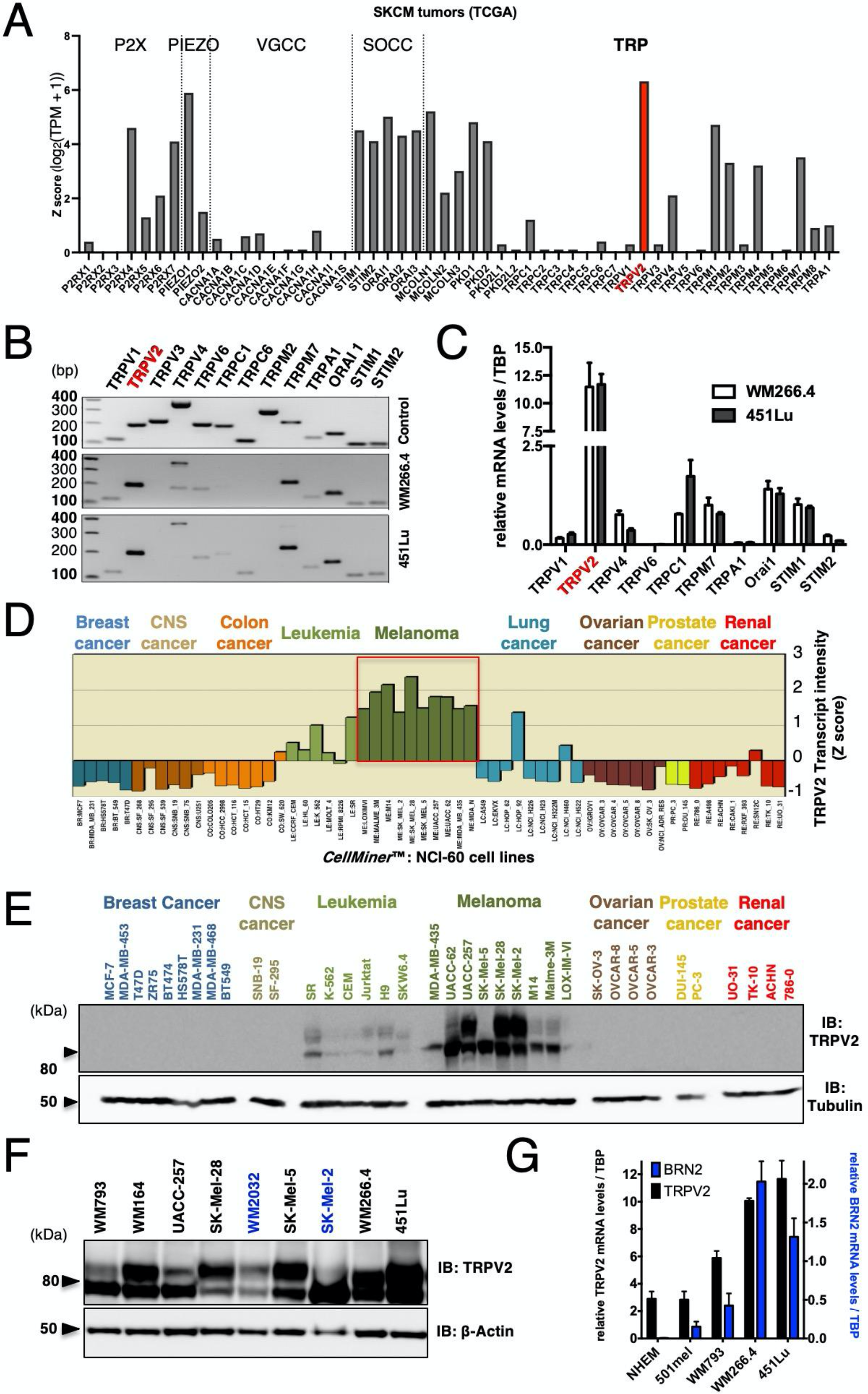
TRPV2 is the predominantly expressed Ca^2+^ channel in metastatic melanoma. (A) Analysis of Ca^2+^-permeable channels mRNA expression levels (from the TCGA cohort RNAseq data) in skin cancer melanoma (SKCM) tumors (n(T)=461). Transcript intensity is expressed as log2(TPM+1) transformed expression data (65). he Ca^2+^-permeable channels plotted on the *x*-axis are grouped by family; ATP-gated P2X; PIEZO; VGCC (voltage-gated Ca^2+^ channels); SOCC (store-operated Ca^2+^ channels); TRP (transient receptor potential). (B) Expression profile of the indicated ion channel analyzed by RT-PCR in the WM266.4 and 451Lu metastatic melanoma cell lines (middle and bottom panels, respectively). Expected amplicons sizes (positive controls in top panel) are in base pairs (bp); TRPV1=120, TRPV2=199, TRPV3=226, TRPV4=190/370, TRPV6=208, TRPC1=201, TRPC6=121, TRPM2=303, TRPM7=226, TRPA1=140, Orai1=161, STIM1=109, STIM2=114). (C) Quantitative RT-PCR expression analyses of channel subunits consistently detected in both the WM266.4 and the 451Lu melanoma cell lines. Transcript levels were normalized to TATA box-binding protein (TBP) mRNA levels. Data are represented as mean ± SEM. (D) Relative TRPV2 mRNA expression analysis in the NCI-60 cell lines panel. *y*-axis represents TRPV2 transcript intensity expressed in mean-centered *z*-score; bars either show increased or decreased expression relative to the mean expression. The cell lines plotted on the *x*-axis are grouped by tissue of origin. CNS: central nervous system. Data were generated by querying for TRPV2 as input in CellMiner^TM^ (http://discover.nci.nih.gov/cellminer/) (66) (See also Figure S1). (E) TRPV2 immunoblotting in 36 out of the 60 NCI-60 cell lines. Tubulin was used as a loading control. Note that after longer exposure time, TRPV2 expression was also detected in some breast and prostate cancer cell lines, as it has been previously described (23,24,67). (F) TRPV2 protein expression in melanoma cell lines harboring either B-RAF (black) or N-Ras (blue) mutations. β-Actin was used as a loading control. (G) Quantitative RT-PCR analysis of both TRPV2 (black bars, left axis) and BRN2 (blue bars, right axis) transcripts expression (normalized to TBP), in normal human epidermal melanocytes (NHEM), non-invasive 501mel, superficial spreading melanoma WM793, metastatic melanoma WM266.4 and 451Lu cell lines. BRN2 is a marker of the melanoma invasive phenotype (see also Figure S6 for active β-catenin levels representing an alternative invasive marker). Data are represented as mean ± SEM.

Molecularly, melanomas harbor somatic ‘driver mutations’ that are mutually exclusive: 50% present gain-of-function BRAF mutations, while another 25% exhibit NRAS mutations (17). By analyzing TRPV2 protein expression in an extended panel of melanoma cell lines harboring either mutation, we consistently observed elevated TRPV2 levels regardless of the mutational status (Figure 1F). Considering its predominant and ubiquitous expression in melanoma, we investigated the functional relevance of this Ca^2+^ channel subunit in melanoma progression.

### TRPV2 expression levels correlate with the invasive phenotype of melanoma tumor cell lines

Melanoma malignancy is mostly driven by its unique ability to rapidly disseminate and form distant metastasis. We therefore investigate TRPV2 expression *versus* melanoma invasiveness, by quantifying TRPV2 transcripts levels along with BRN2 transcription factor mRNAs, a well-established invasive phenotype marker (18). While very low levels of TRPV2 mRNAs were present in normal human epithelial melanocytes (NHEM), a gradual increase of TRPV2 transcripts mimicked the rise of BRN2 expression in melanoma cells (ranging from the non-invasive 501mel, to the superficial spreading melanoma WM793, then to the metastatic melanoma WM266.4 and 451Lu) (Figure 1G). This positive correlation between invasiveness and TRPV2 expression was further confirmed at the protein level (Figure 2A) and by a functional approach (Figure S2A). TRPV2 channel over-activation by its potent agonist cannabidiol (CBD) (19) induced a characteristic intracellular [Ca^2+^] increase, where the extent of functional channels matched with the total amount of proteins, ultimately connecting the expression of functional TRPV2 channels to the invasive phenotype of melanoma cell lines.

**Figure 2.**
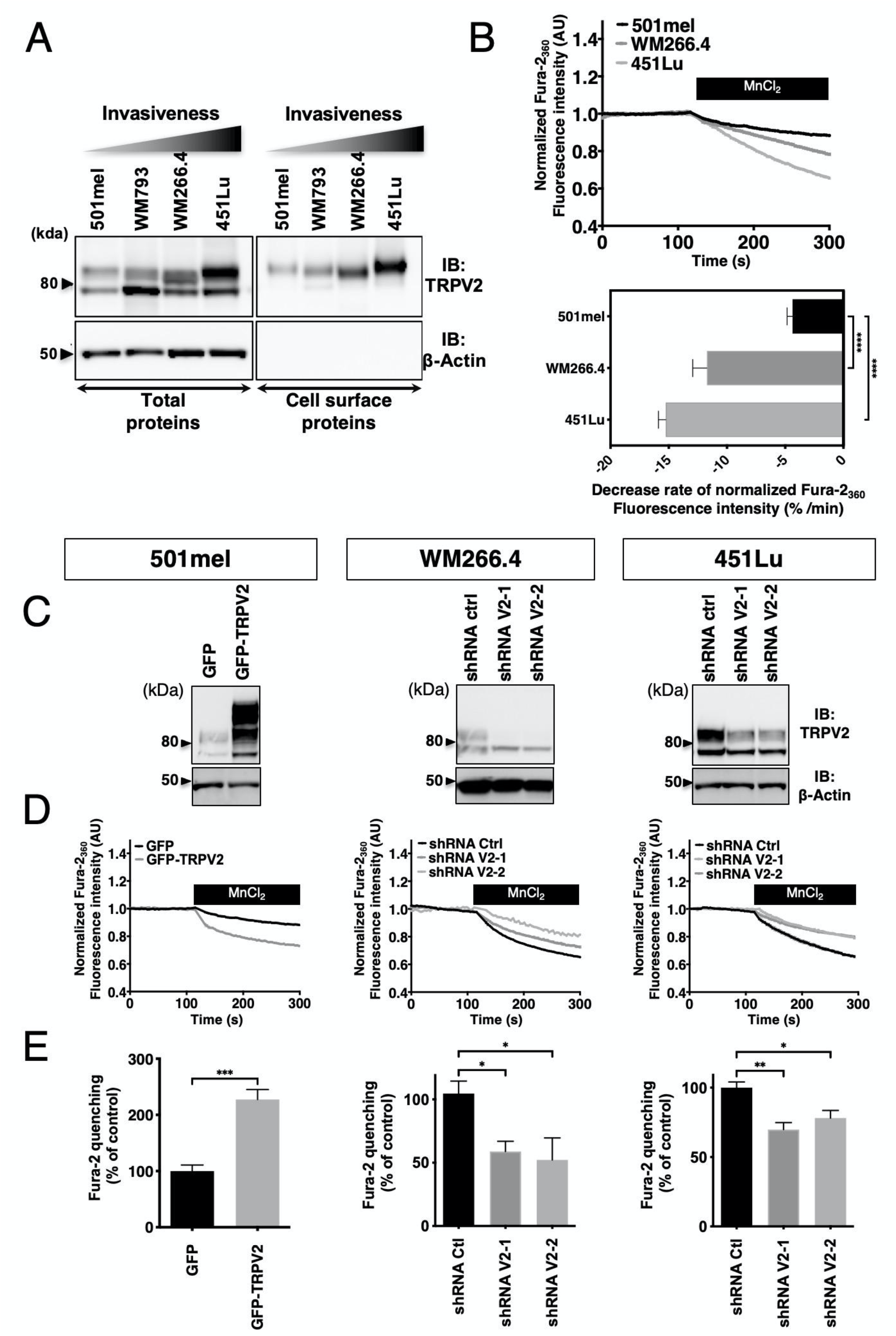
TRPV2 expression, plasma membrane targeting and contribution to basal Ca^2+^ influx, increase with melanoma cell aggressiveness. (A) Analysis for total and plasma membrane (PM) TRPV2 expression in melanoma cell lines as in A. β-actin was used as loading and cell integrity control. PM fractions were isolated using the surface protein biotinylation technique. The higher molecular weight bands of TRPV2 correspond to the glycosylated and mature channel (50) (See also Figure S2). (B) Constitutive Ca^2+^ influx comparison between resting 501mel (black), WM266.4 (dark grey) and 451Lu cells (light grey) assessed using the Fura-2 Mn^2+^ quenching assay. Representative traces of Fura-2 quenching kinetics are shown on the top panel (each point represents the mean of quadruplicates normalized to the baseline). The bar graph summarizes the quantification of the measured quenching rates and is presented as mean ± SEM. (C) TRPV2 overexpression in the non-invasive 501mel cell line (GFP-TRPV2) or downregulation by lentiviral-delivery of TRPV2 specific shRNAs (shRNA V2-1 and V2-2) in the WM266.4 and 451Lu metastatic melanoma cell lines assessed by western-blot. β-actin was used as a loading control. (D) Representative traces of Fura-2 Mn^2+^ quenching rates in control (GFP) or GFP-TRPV2 overexpressing 501mel cells, or in control (shRNA Ctrl) or TRPV2 repressed WM266.4 and 451Lu cells. (E) Average normalized quenching rates (as in D) presented as mean ± SEM.

### TRPV2 channels are addressed to the plasma membrane and active in metastatic melanoma cells

Plasma membrane (PM) trafficking has been postulated as an important regulatory mechanism for TRPV2 activity (20). Surface protein biotinylation experiments evidenced that the subset of TRPV2 channels present at the PM increased gradually with the invasive potential of the tested melanoma cell lines (Figure 2A). When assessed by confocal immunofluorescence, TRPV2 was consistently detected at the PM of the highly metastatic melanoma cell lines, WM266.4 and 451Lu (Figure S2B). The measurement of non-stimulated Ca^2+^ influx with the Mn^2+^ quenching assay next correlated the amplitude of the constitutive Ca^2+^ entry to the extent of TRPV2 distribution at the PM. Indeed, in the 501mel cells, where sparse TRPV2 labelling mostly remained intracellular, the quenching rate was weak as opposed to both metastatic cell lines harboring PM-resident TRPV2, with the 451Lu cells displaying the highest constitutive Ca^2+^ entry (Figure 2B).

To determine whether resting Ca^2+^ entries directly depend upon TRPV2 function in advanced melanoma, we modulated TRPV2 expression in three cell lines with well-defined, but opposite invasive phenotypes (Figures 1G, S6 and (35)). TRPV2 was either over-expressed in the non-invasive 501mel cell line, or silenced with two different shRNA sequences in the highly invasive WM266.4 and 451Lu cell lines. Successful TRPV2 overexpression or repression was confirmed by immunoblotting (Figure 2C), and Ca^2+^ imaging experiments further validated the channel functionality (Figure S2C). GFP-TRPV2 overexpression in 501mel cells yielded an elevated response to CBD, whereas TRPV2 silencing (shRNA-V2) in WM266.4 and 451Lu cells, by disabling protein expression, hampered CBD-induced Ca^2+^ influx. Resting Ca^2+^ signals in non-stimulated adherent cells then showed that TRPV2 overexpression in 501mel cells doubled the basal Ca^2+^-influx, while TRPV2 silencing in WM266.4 and 451Lu cells weakened it (Figures 2D-E). In unstimulated metastatic melanoma cells, a subset of TRPV2 channels is therefore addressed to the PM and is active, allowing Ca^2+^ entry.

### TRPV2 is dispensable for cell proliferation but is essential for melanoma tumor cell migration and invasion

Depending on the tumoral context, TRPV2 has been shown to be specifically implicated in proliferation, and/or in the progression toward a pro-invasive phenotype (21). We therefore evaluated whether TRPV2 expression affects either of these hallmarks in melanoma. Although, TRPV2 overexpression increased the growth rate of the non-invasive 501mel cells, TRPV2 silencing had no impact on the viability nor on ERK phosphorylation in both metastatic cell lines, suggesting that TRPV2 is dispensable for malignant melanoma proliferative/survival behavior (Figures S3A-B). However, upon TRPV2 silencing in the metastatic melanoma cell lines (WM266.4 and 451Lu), both serum-induced migration and, to a greater extent, invasion (through a matrigel layer) were strongly hampered (by 40-80% and 65-95%, respectively) (Figure 3A). Reciprocally, TRPV2 overexpression in the 501mel cells was sufficient to increase both their migratory and invasive capacities by 2.5 and 1.3 folds respectively, revealing TRPV2 expression as determinant for melanoma cells migration and invasion potentials.

**Figure 3.**
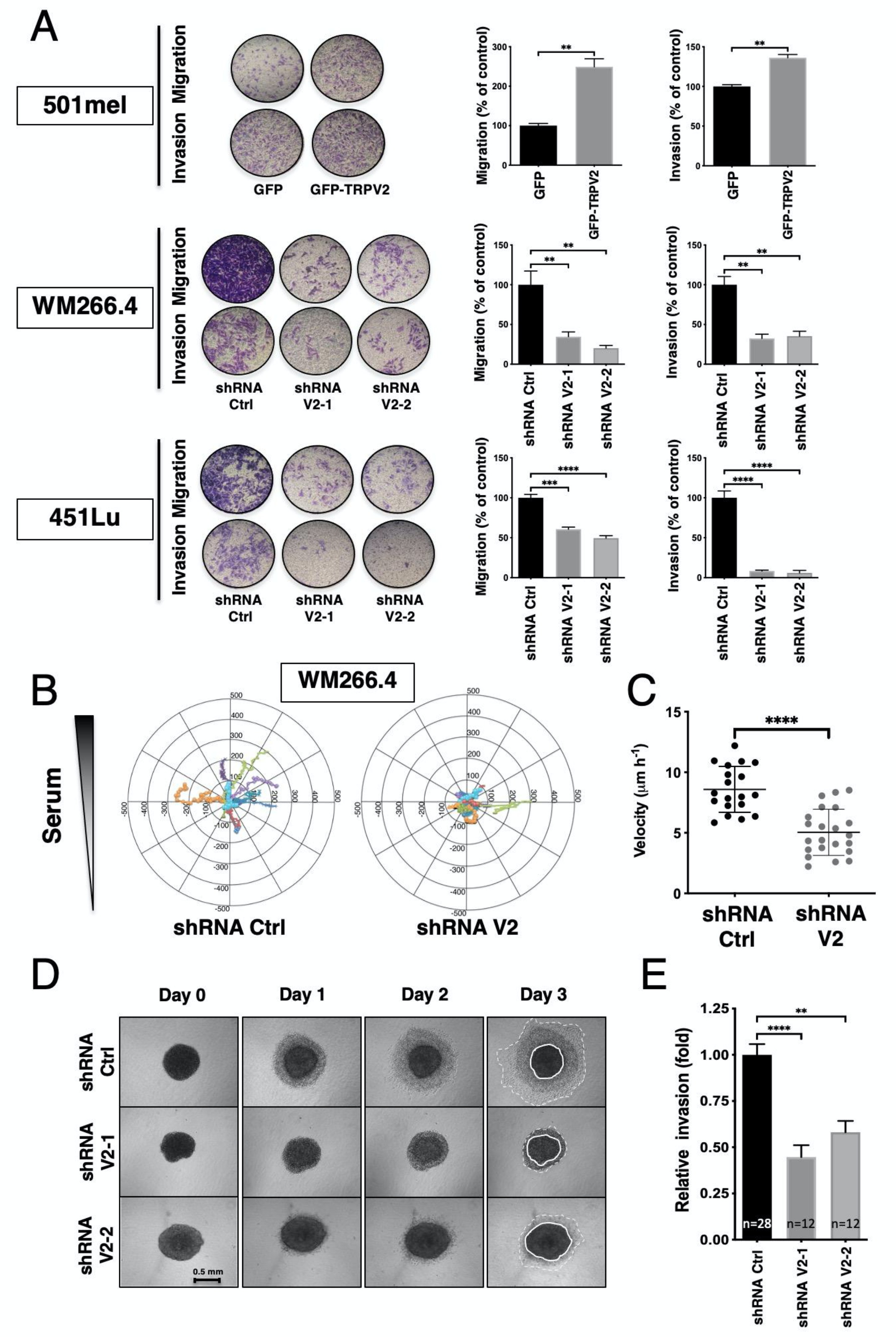
TRPV2 is essential for melanoma tumor cell migration and invasion. (A) Impact of TRPV2 genetic manipulation in 501mel, WM266.4 and 451Lu on serum-induced migration and matrigel invasion. Representative pictures show crystal violet-stained cells that have migrated through 8 µm pore size membranes. Histograms illustrate the average numbers of migrating/invading cells normalized to control (presented as mean ± SEM) from three independent experiments. (B) Tracks comparison between fibronectin-plated shRNA Ctrl or shRNA V2 WM266.4 cells migrating toward a serum gradient over a 12 h period (n=10 and 12, respectively) (See also Figure S4A). (C) Velocity analysis of 2D migration experiments described in B (dot, single cell, n=19-21). Scatter plots show mean ± SD. (D) TRPV2-silencing effect on 3D invasion. Representative images were taken every day for 3 days after collagen embedding of spheroids from WM266.4 cells expressing either shRNA ctrl or TRPV2-targeting shRNAs (scale bar=0.5 mm). (E) Quantification of collagen invasion. For each spheroid, the cell-covered area at day 2 was normalized to the starting area of the collagen-embedded spheroids. Histograms represent the invasion relative to control WM266.4 spheroids (shRNA Ctrl) from at least 12 spheroids from three independent experiments (See also Figure S4B). Data are presented as mean ± SEM.

A detailed motility analysis of the WM266.4 cells, presenting well-adapted morphological characteristics for individual cell 2D-tracking, was carried out in the presence of a serum gradient (Figure S4A). Differentially labeled shRNA-Ctrl and -V2 expressing cells were mixed, seeded and recorded simultaneously (Supplemental Movie 1). Although cell spreading and lamellipodial protrusions appeared as constant features, motility was considerably altered in TRPV2-silenced cells, as attested by a robust reduction of speed and random displacement, compared to control cells (Figures 3B-C). Additionally, to recapitulate the *in situ* tumor-confined environment, WM266.4 cells were grown as melanospheres, embedded in collagen-I matrices, and 3D dynamics was followed for 3 days. Corroborating our 2D-data, TRPV2 repression drastically precluded metastatic melanoma cell 3D-invasion capacity (Figures 3D-E and S4B). Interestingly, we did not observe any significant difference in the size of the melanospheres formed by the WM266.4 cells, whether expressing TRPV2 or not (not shown), confirming that TRPV2 silencing does not disrupt advanced melanoma cells proliferation or survival potentials.

Migratory melanoma cells display high plasticity, and their migration could result from a combination of several interconnected mechanisms, notably among a pseudo-epithelial–mesenchymal transition (pseudo-EMT) and interactions with the microenvironment. When we assessed EMT-associated markers, 501Mel and WM266.4 cells exhibited distinctively opposite profiles, epithelial- or mesenchymal-like respectively, while 451Lu cells presented an intermediate phenotype. Most importantly, neither TRPV2 overexpression nor its down-regulation altered the levels of pseudo-EMT markers (Figure S6).

### TRPV2 associates with nascent acto-adhesive structures in metastatic melanoma cells

Cell migration requires highly coordinated interactions between the extracellular matrix and the intracellular cytoskeleton *via* multiprotein adhesion structures (22). These structures dynamically transition upon mechanical tension from nascent adhesions to focal complexes and focal adhesions (FAs) (23). In spite of a critical role established for Ca^2+^ signaling in both adhesion and actin cytoskeleton dynamics (24, 25), no studies in cancer cells inferred a direct function for the mechanosensitive channel TRPV2 in these processes. However, a large-scale proteomic analysis detected TRPV2 as the only Ca^2+^ channel present in adhesion structures (26), prompting us to explore its potential interaction with the acto-adhesive machinery in the dynamic context of migrative melanoma cells. In migratory metastatic WM266.4 and 451Lu cells, PM endogenous TRPV2 was frequently distributed at the leading edge of the cells and, in many cases a sharp staining was found near the lamella. In these define proximal clusters, TRPV2 signal preferentially colocalized with the early phase marker of adhesion, paxillin, and with F-actin, rather than with the markers of mechanically engaged FAs, vinculin or activated pY397-FAK (Figures 4A-B) (23). A combination of proximity-ligation assays and spatial distribution analysis of molecules by dual-color direct stochastic optical reconstruction microscopy (dSTORM), clearly attested the preferential co-clustering of TRPV2 channels with paxillin-rather than vinculin-containing structures in migrating metastatic melanoma cells (Figures 4C-D and S5A). Additionally, we noted that TRPV2 co-localized and physically interacts with vimentin (Figures S5B-C), further arguing for TRPV2 channels involvement in the mechanical regulation of nascent adhesions (27, 28).

**Figure 4.**
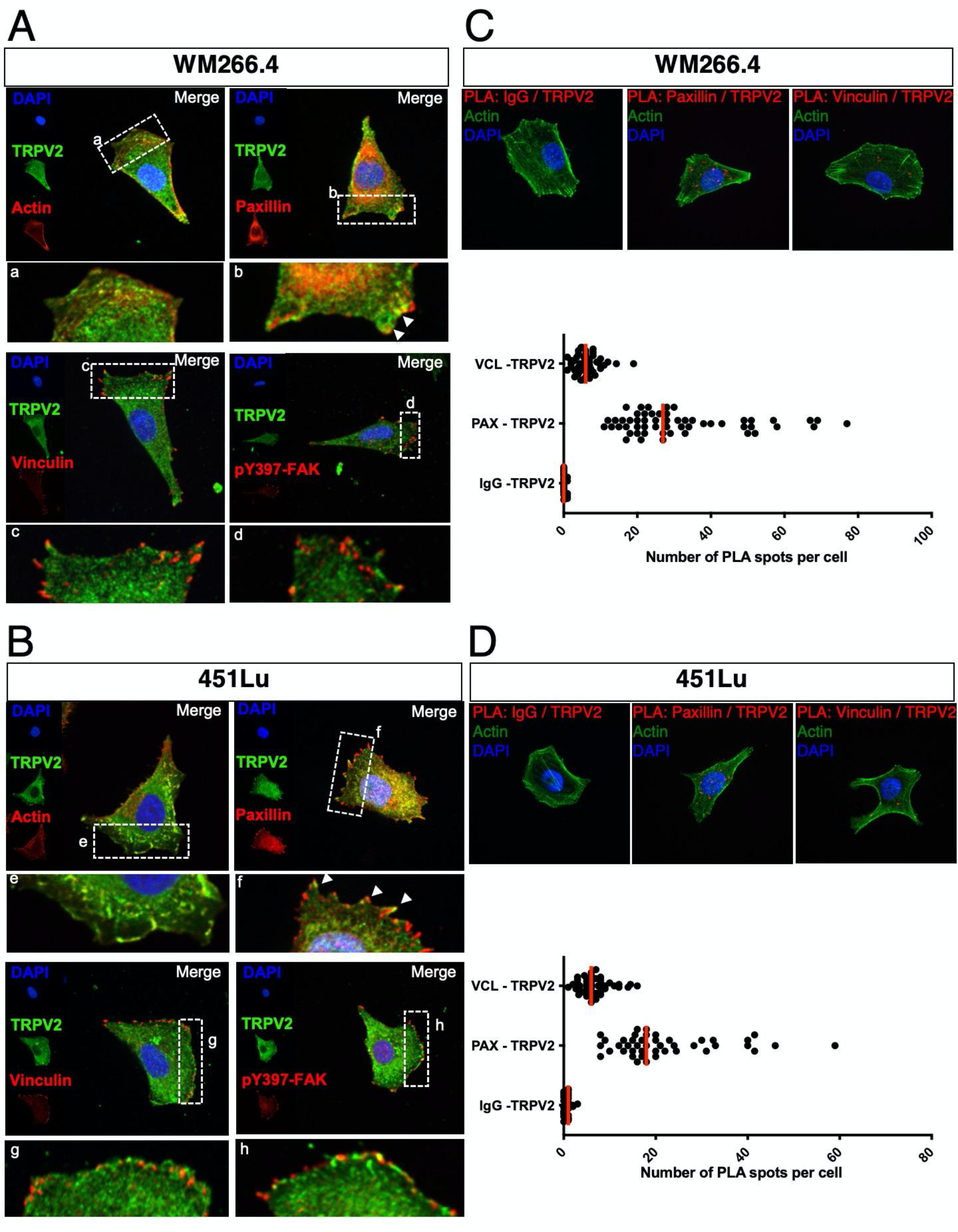
TRPV2 associates with markers of nascent acto-adhesive structures. Representative confocal images of low confluency WM266.4 (A&C) and 451Lu (B&D) metastatic melanoma cells seeded on fibronectin-coated coverslips. (A & B) Cell nuclei are depicted with DAPI in blue, TRPV2 in green and indicated protein (actin, paxillin, vinculin or pY397-FAK) in red. Insets are magnifications of the indicated area. Arrows indicate sites of colocalization. (C & D) F-actin staining (green), cell nuclei (blue) and proximity-ligation assay (PLA) reaction between indicated proteins (red). Red fluorescent spots indicate the association of the two proteins of interest, close to 40 nm. Scatter plots represent the quantification of the number of PLA spots per cell (bars indicate the medians) between TRPV2 and a control antibody (IgG), paxillin (PAX) or vinculin (VCL) (See also Figure S5).

### TRPV2 modulates melanoma tumor cell migration through the control of the calpain-dependent maturation of FAs and actin cytoskeleton remodeling

To investigate how TRPV2 participates in adhesion complex dynamics, we evaluated TRPV2 impact over FAs maturation. Mechanically engaged adhesions were detected and quantified following vinculin staining (Figures 5A and S7A). Overexpressing TRPV2 in the 501mel cell line reduced by 23% the occurrence of FAs, while TRPV2 silencing in the WM266.4 and 451Lu cells increased the frequency of FAs by 27% and 24%, respectively. Of note, modulating TRPV2 expression either way did not impact total vinculin levels (Figure S7B). A known Ca^2+^-dependent mechanism for adhesion disassembly involved the calpain-mediated proteolysis of talin, a mechanosensitive adhesion protein described as interfering with vinculin recruitment and FAs mechanical engagement (29–31). We therefore tested whether TRPV2 could regulate the activity of the Ca^2+^-activated calpains. In the 501mel cell line, TRPV2 overexpression induced a 2-fold increase in calpain basal activity compared to MOCK cells, reciprocally TRPV2 silencing in both metastatic melanoma cell lines halved the protease activity (Figure 5B). Regarding talin proteolytic degradation, which largely depends upon extracellular Ca^2+^ signaling in our melanoma models (Figures S7C-D), it exactly mirrored TRPV2-mediated calpain activity. Talin cleaved isoform (190 kDa) increased by 75% in 501mel cells overexpressing TRPV2 compared to control cells mostly exhibiting full-length form (230 kDa), evoking FAs stability and the low motility of these cells (Figure 5C). In the malignant WM266.4 and 451Lu cells, the cleaved-talin isoform was distinctly detected, which again corroborates with adhesion plasticity in these highly migrating cells. Upon TRPV2 silencing calpain-mediated cleavage of talin decreased by 60 to 75%. Altogether we showed that TRPV2 directly regulates calpain activation and the subsequent cleavage of talin, to control adhesion dynamics.

**Figure 5.**
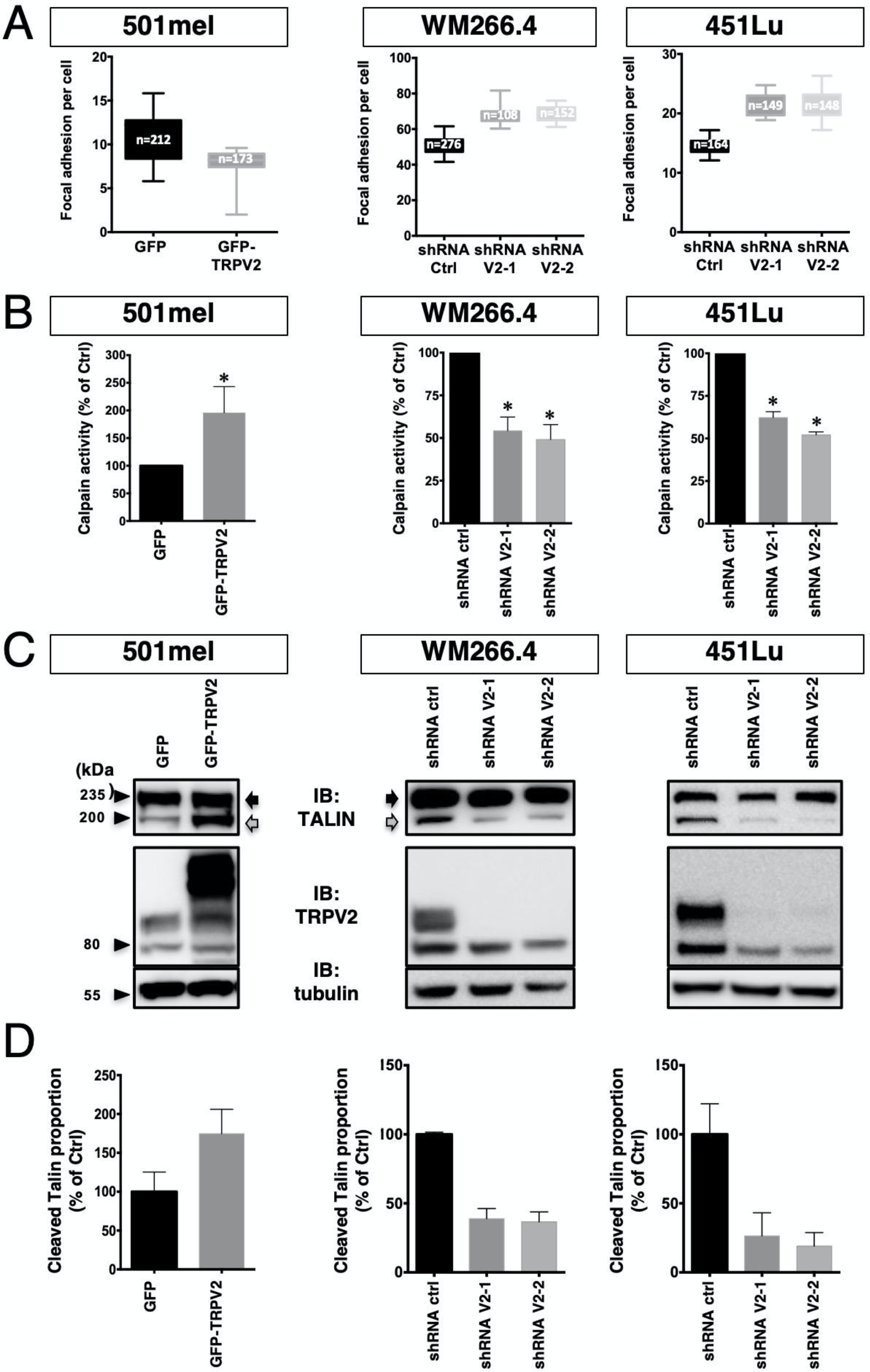
TRPV2 activity controls the mechanical maturation of focal adhesions. (A) Quantification of vinculin-stained focal adhesion sites per cell. Images were analyzed using the imageJ software to count the number of vinculin clusters per cell. The number of cells counted for each cell line is indicated in the boxes. Data are presented as min-to-max whisker-box plots. (B) Histograms represent normalized calpain activity in control (GFP) or overexpressing TRPV2 (GFP-TRPV2) 501mel cells, and in control (shRNA Ctrl) or TRPV2-silenced (shRNA V2-1, −2) WM266.4 cells or 451Lu cells. Bar graphs show mean ± SEM (n=3 independent experiments). (C) Representative immunoblots showing the full-lenght (230 kDa) and the calpain-mediated degradation product (190 kDa) of talin after overexpression or repression of TRPV2, in the corresponding cell lines. Tubulin was used as a loading control. (D) Densitometric analysis of the cleaved-talin ratios from 3 independent experiments as described in C normalized to control. Data are presented as mean ± SEM.

FAs growth cooperatively reinforces the link between the ECM and cytoskeleton, with adhesion components, such as talin, directly being at this interface. Since cellular migration equally depends on cytoskeletal remodeling, and TRPV2 activation has been related to mechanically-induced actin assembly/disassembly rate (32), we analyzed filamentous actin (F-actin) cytoskeleton, quantified F-actin bundles and normalized to the cell area in migrating CMM models (Figures S8A-B). Despite a globally steady melanoma cells area, TRPV2 overexpression in 501mel cells was by itself sufficient to considerably promote F-actin accumulation, whereas TRPV2 silencing in aggressive melanoma cells disrupted F-actin network. In a previous proteomic screen, we have identified cofilin-1, known modulator of F-actin dynamics, as a potential TRPV2-interacting protein (unpublished data). Here, we showed that TRPV2 physically interacts with cofilin-1 in metastatic melanoma cell lines (Figure S8C). In WM266.4 cells and, to a lesser extent in 451Lu cells, we observed that TRPV2 silencing reduced the level of inactive cofilin-1 (phosphorylated form), suggesting that TRPV2 represses cofilin-1 activity (Figure S8D). Taken together, our results highlight a new role for TRPV2 in regulating advanced melanoma cell motility through the control of calpain-mediated FAs mechanical maturation conjointly to cofilin-1-induced actin cytoskeleton reorganization.

### TRPV2 expression level determines the *in vivo* metastatic potential of melanoma tumor cells

To determine whether TRPV2 activity ultimately impacts the formation of metastasis *in vivo*, we injected bioluminescent human melanoma cells displaying modulated expression levels of TRPV2 into the tail vein of immunocompromised mice, and followed metastasis formation by bioluminescence imaging (BLI). At the assay end point, *in vivo* BLI showed a tendency towards an increasing metastatic potential for GFP-TRPV2 overexpressing 501mel cells (Figure 6A). Unequivocally, *ex vivo* BLI at necropsy uncovered a dramatic increase of the metastatic burden in mice injected with GFP-TRPV2 overexpressing cells compared to controls. Numerous metastatic foci were observed in lungs, brain and bones (Figures 6B and S9A-B), revealing that TRPV2 expression was sufficient to endow 501mel cells with metastatic competences. In line with this observation, TRPV2 silencing in 451Lu cells significantly decreased their metastatic potential in mice. BLI immediately post-injection revealed that TRPV2 repression prevented the extravasation of melanoma cells into the lungs (Figure S9C). At the assay end point, both *in vivo* and ex *vivo* BLI further confirmed that TRPV2-depleted cells have lost their potential in long-term lung colonization compared to control cells (Figures 6C-D). Immunofluorescence analysis in lung sections eventually showed that individual metastatic lesions in control-injected mice retained endogenous TRPV2 expression, strongly detected at the cell periphery (Figure S10). TRPV2 requirement for the formation of melanoma metastasis was likewise validated in the experimental model of xenografted zebrafish allowing a direct comparison of the metastatic potential of two different cell lines in the same organism. To distinguish between control- and TRPV2-shRNA WM266.4 expressing cells that are both GFP-labeled, control cells were double-stained with the red fluorescent dye CmDil. Then, equal amounts of both shRNA-expressing cells were mixed and co-injected in the duct of Cuvier of 2 days-old zebrafish embryos. Thirty-six hours post-transplantation, only double-stained control cells, expressing TRPV2, have disseminated throughout the fish body (Figure S11). Altogether, these results established that melanoma tumor cells rely on TRPV2 to succeed in disseminating and form distant metastases.

**Figure 6.**
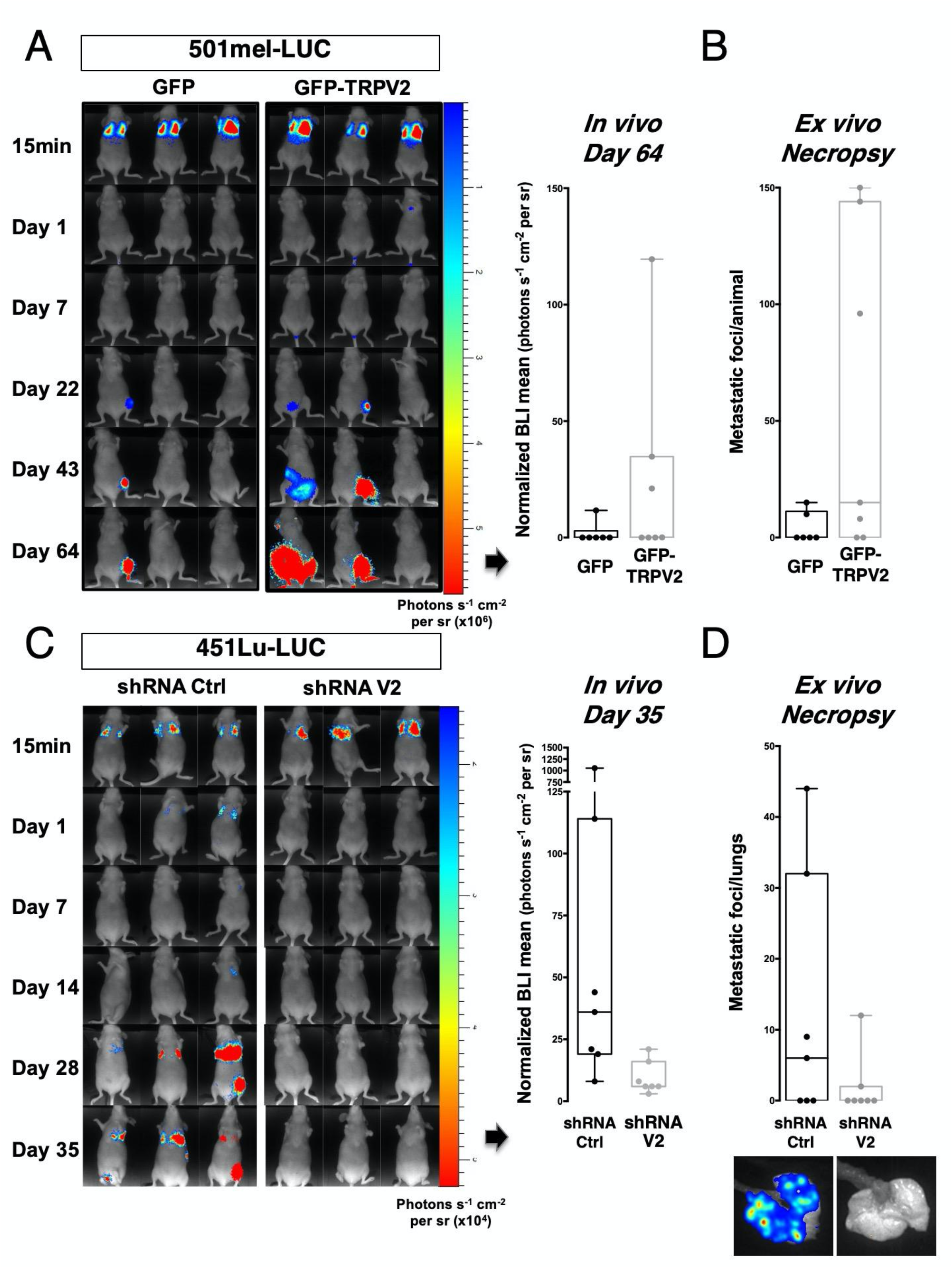
TRPV2 expression level determines the *in vivo* metastatic potential of melanoma tumor cells. (A & C) Representative bioluminescence imaging (BLI) data of mice injected intravenously with the non-metastatic melanoma cell line 501mel-Luc transfected with either GFP-TRPV2 or GFP control (A), or with the invasive 451Lu-Luc melanoma cell line expressing either control shRNA or TRPV2 targeting shRNA (C). Tumor growth and metastasis formation were monitored for 64 days (501mel-Luc) or 35 days (451Lu-Luc) after injection. Normalized photon flux quantification at the end time point is represented (dot, single mouse, n=6-7). Data are presented as min-to-max whisker-box plots (See also Figure S9-10). (B & D) Scatter plot of the number of metastatic foci per animal (B) or per lungs (D) counted at necropsy of either (B) mice xenografted with GFP control (black) or GFP-TRPV2 overexpressing (gray) 501mel-Luc cells (dot, single mouse, n=6-7), or (D) mice injected with shRNA Ctrl (black) or shRNA TRPV2 (gray) 451Lu-Luc cells (dot, single mouse, n=7). Representative *ex vivo* BLI of lung metastasis is shown (bottom). Data are presented as min-to-max whisker-box plots.

### TRPV2 expression in melanoma as a marker of advanced malignancy and bad prognosis

The interdependence between TRPV2 and the metastatic phenotype of melanoma tumor cells *in vitro* and *in vivo* prompted us to evaluate the clinical significance of TRPV2 expression in human melanoma. The TCGA skin melanoma RNAseq dataset was first compared to the matched TCGA and GTEx normal datasets (Figure S13A). Consistent with NHEM *versus* melanoma cell lines observations (Figure 2A), TRPV2 mRNA level was significantly higher in melanoma compared to healthy skin melanocytes. *In situ* protein expression was then assessed by combining immunohistochemical and tissue microarray (TMA) analyses (Figures 7A-C and S12A-B). The TMA assembled 100 patient samples including 62 malignant melanomas (Grade I-IV), 20 lymph node metastases and 18 benign nevi tissues (Figure S12C). This tissue cohort analysis demonstrated that overall malignant melanomas and lymph node metastasis were extensively expressing TRPV2, while benign nevi exhibited no or weak staining (Figures 7A-C and S12D). With regards to melanoma progression, TRPV2 intensity appeared correlated with advanced lesions (Figures 7B-C and S13B), albeit statistical significance between malignant *versus* metastatic tissues was not reached, likely due to a limited number of high-grade samples (4 grade III and 2 grade IV). Yet, a similar trend was observed on randomly selected stages I and IV melanomas issued from a local hospital independent tumor biobank (Figure S13C). Moreover, analysis of TRPV2 transcript level according to the Clark level staging system, defining anatomical invasion, revealed that higher the expression of TRPV2 was, the deeper the tumor had penetrated into the skin layers (Figure 7D). Comparison of isogenic cell lines pairs, such as WM164/451Lu or WM115/WM266.4, further established that TRPV2 expression was higher in metastasis derived cell lines compared to the poorly invasive matched cell lines originated from *in situ* tumors (Figures 1F and 7E). Finally, survival analysis evidenced that melanoma tumors expressing high levels of TRPV2 correlated with shorter life expectancies in patients, compared to low TRPV2 expressers (Figure 7F).

**Figure 7.**
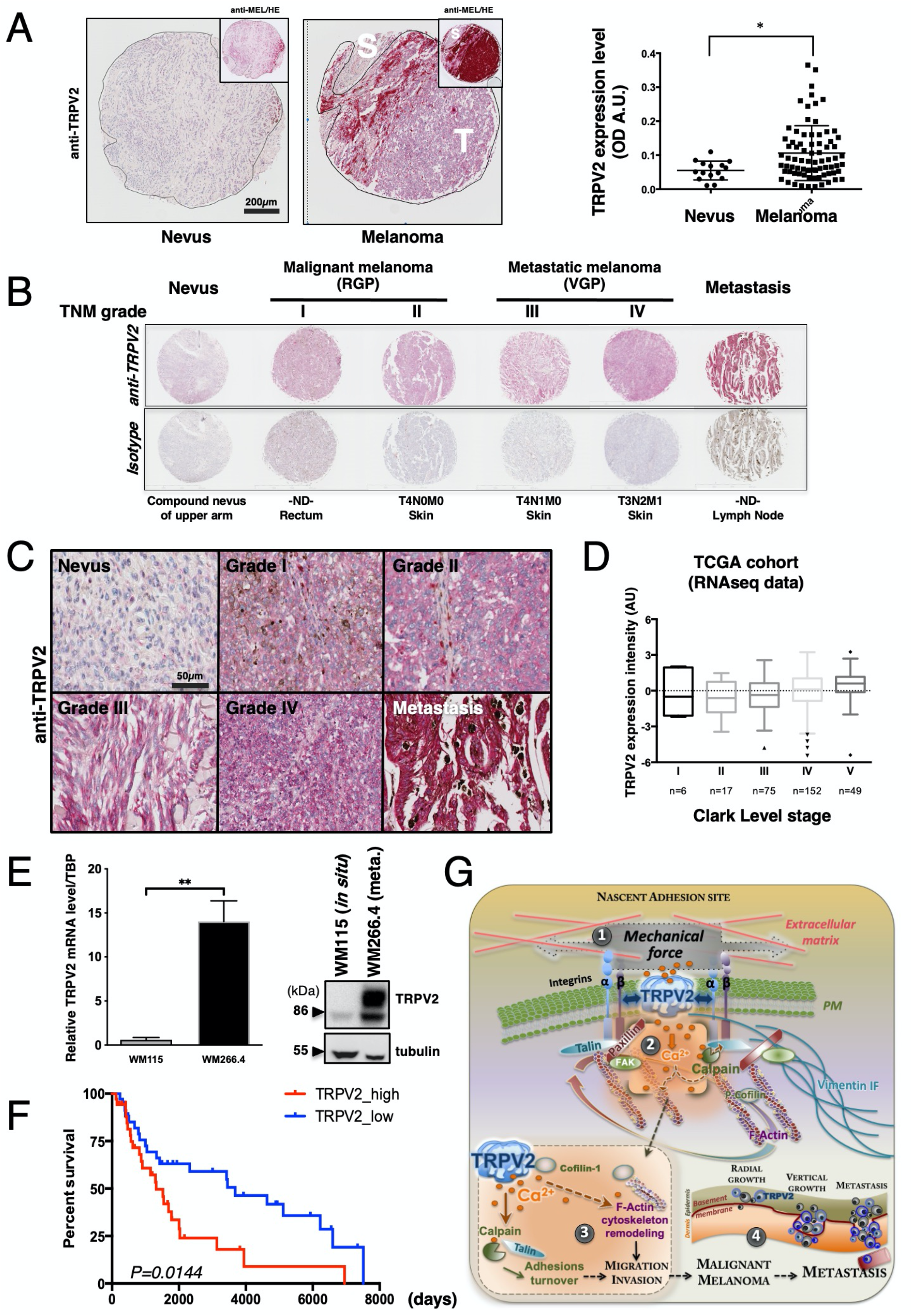
TRPV2 overexpression in melanoma is a marker of advanced malignancy and bad prognosis. (A) Representative melanoma tissue lesions from a tissue microarray (TMA) comparing TRPV2 staining between a benign nevus and a malignant melanoma. Top right insets show the same images stained with an anti-melanoma triple cocktail (HMB45+A103+T311). Regions depicted with a black line represent the interface between the diffusely stained tumor (T) and the surrounding normal stroma (S). Right panel shows the scatter plot of the complete quantification of TRPV2 staining in 82 patients with malignant melanoma compared to 18 nevi tissues (dot, single patient sample) Scatter plots show mean ± SEM. (See also Figure S12). (B) Comparison of 6 representative tissues from patients at different TNM grades of melanoma progression stained with anti-TRPV2 or the isotype control. RGP: radial growth phase, VGP: vertical growth phase. (C) Representative images showing detailed TRPV2 staining from *in situ* melanoma lesions with progressive TNM grades and from lymph node (LN) metastasis. (D) TRPV2 mRNA expression (from the TCGA cohort RNAseq data) according to Clark level pathological cancer stages. This grading system describes the level of anatomical invasion of the melanoma; I: confined to the epidermis (*in situ*); II: invasion into the papillary dermis; III: invasion to the junction of the papillary and reticular dermis; IV: invasion into the reticular dermis; V: invasion into the subcutaneous fat. Data is represented as a whisker-box plot and outliers are plotted as individual points (n are indicated on the plot) (See also Figure S13). (E) Analysis of TRPV2 expression by quantitative RT-PCR (left panel) and western-blot (right panel) in the WM115/WM266.4 pair of isogenic melanoma cell lines. The WM115 cell line was derived from an *in situ* tumor while the WM266.4 was established from a skin metastasis of the same patient. Relative transcript levels are presented as mean ± SEM. (F) Kaplan–Meier plot showing the association of TRPV2 expression (10% highest *versus* 10% lowest expressers in the skin cutaneous melanoma cohort TCGA data set) with melanoma patient survival. (G) Mechanistic model of melanoma tumor cell dynamics regulation by TRPV2. The essential role of TRPV2 in melanoma migration and invasion could be explained by the newly identified pro-invasive properties of this mechanosensitive channel. Indeed, cancer cells and their associated microenvironment generate considerable mechanical forces applied onto the plasma membrane (PM) (1). These changes in PM tension regulate cell shape and movement. In malignant melanoma cells, the mechanosensitive TRPV2 channel is recruited at the PM within paxillin-containing early adhesion structures, and its constitutive and/or mechano-activation yield to a subplasmalemmal localized Ca^2+^ ions uptake (2). TRPV2-mediated Ca^2+^ influx triggers the activation of the intracellular Ca^2+^-dependent cysteine protease, calpain (3). The cleavage of its substrate, the adhesion protein talin linking membrane integrins and cytoskeleton, *in fine* triggers some adhesion complexes disassembly and facilitates cell-extracellular matrix (ECM) contact sites plasticity. Induction, selection and maturation of nascent adhesion complexes at the cell leading edge serve as sampling the local ECM to select traction points producing forces that will drive the cell body forward. To further regulate the maturation of these adhesion structures along with the remodelling of the cytoskeleton, TRPV2 directly interacts with both the intermediate filament (IF) vimentin network, and the actin severing factor cofilin-1, a central regulator of F-actin dynamics. TRPV2-induced signaling promotes the spatial and temporal accumulation of F-actin bundles to improve advanced melanoma cell motility. Therefore, TRPV2 channel-mediated Ca^2+^ influx tunes the plasticity of the melanoma tumor cell by directly and locally controlling adhesion complexes maturation and cytoskeleton remodeling, potentiating the migratory and invasive behavior of these malignant cells (4).

## DISCUSSION

Despite being one of the least characterized TRP members (33), potential roles for TRPV2 have been evoked in various cancers during the last decade (21). Here, we established that in melanoma tumor cells the expression of functional TRPV2 channels was associated with invasiveness, making TRPV2 mandatory for the dissemination and formation of distant metastases. Similarly, in melanoma biopsies, TRPV2 expression increased together with tumor progression, invasive phenotype, metastatic potential and ultimately a shorter life expectancy. Consistent with reports based on tumors from different origins, including esophageal squamous cell, breast, prostate and urothelial carcinomas, as well as glioblastoma (34–37), our results set TRPV2 as a valuable prognosis marker in melanoma tumor progression.

TRPV2 expression levels have been found deregulated in patients with metastatic disease compared to primary solid tumors (21). Among the biological processes regulated by TRPV2 channel during metastatic progression, TRPV2 has been notably associated with proliferation, for instance in esophageal squamous cell carcinoma (35), or with the progression toward a pro-invasive phenotype, in prostate and bladder cancers (34,38,39). Melanoma is an aggressive cancer endowed with unique features of cellular plasticity, coupled to a rare ability to switch back and forth between proliferative and invasive phenotypes. Using gain- and loss-of-function approaches, we established that TRPV2 expression and activity potentiates the acquisition of both the migratory and invasive phenotypes of melanoma cells, while dispensable for their proliferative/survival behaviors. Note that specifically addressing TRPV2 role, solely by modulating its expression, prevents the issue of indirect/off-target Ca^2+^ effects often raised with pharmaco-modulators and maybe accounting for some disparities seen in the literature.

We investigated whether the mechanistic basis of TRPV2-mediated aggressive potential in metastatic melanoma cells, relied on a pseudo-EMT program which in some cancer cells is regulated by Ca^2+^ signaling (40, 41). Note that for melanoma the term EMT remains inadequate since melanocytes are not epithelial cells and their invasive state may not be mesenchymal. Nevertheless, modulating TRPV2 expression - either way - had no impact on EMT markers’ levels. We however noted that the non-invasive 501Mel *versus* the highly invasive WM266.4 cells were exhibiting antagonistic markers profiles, corresponding to either a pseudo-epithelial or mesenchymal-associated state respectively, and suggesting that the WM266.4 have completed a transition process to acquire their metastatic potential. These observations were in adequation with their morphology and migrative features, as well as with their levels of BRN2 invasiveness marker. Regarding the 451Lu cells, the expression profiles of pseudo-EMT and BRN2 markers were representative of a previously described “intermediate phenotype” (42). These cells in a “partial” state defined as capable of both adhesion and migration, are matching with the 451Lu extreme invasive potential associated with an amoeboid/mesenchymal morphology and fewer adhesion structures compared to the WM266.4 cells. Altogether reflecting the highly heterogeneous and dynamic properties of metastatic melanoma cells, which have been described as driven by the tumor microenvironment (TME).

The microenvironment is indeed thought to contribute to the process of metastasis, and is clearly of importance for both cell migration and invasion (43). In unstimulated melanoma cells, we evidenced that TRPV2 is localized at the PM and is active, yet we excluded a growth factor-dependent regulation of its trafficking (data not shown). We however know that the dynamic endosomal-PM translocation of TRPV2, regulating its activity (44), can be induced by mechanical cues (5). Hence, adhesion dynamics and the resulting mechanical constraints applied to the PM could be critical factors controlling TRPV2 activity. Therefore, globally impacting resting Ca^2+^ homeostasis which has been directly correlated to melanoma aggressiveness (8). The constant changes in cell-matrix contact points and cell shape are guided by mechanical cues triggering adhesions dynamics and actin cytoskeleton remodeling (45). Soon after the cell-matrix engagement, the adapter protein paxillin gets recruited to form nascent adhesions. To date, TRPV2 is the only Ca^2+^-permeant channel reported as directly participating in paxillin-rich adhesive structures, revealing a mechanism by which TRPV2 could boost melanoma tumor cell invasiveness. Regulations of these nascent adhesion structures may directly influence sensing, forces generation and maturation of adhesion complexes that drive the cell body forward. Generally, paxillin clusters are found at the proximal end of the FA, where paxillin strongly binds to integrins during the early phases of FA formation, and where F-actin also enters and accumulates, defining paxillin as actively engaged in highly dynamic complexes (46). In migrating CMM cells, we also determined that TRPV2 prompted actin filament accumulation together with cofilin-1 inactivation. We showed that TRPV2 physically interacts with cofilin-1, this actin severing factor known to coordinate the spatiotemporal organization of F-actin during cell migration by integrating transmembrane signals (47). Knowing that TRPV2-dependent actin rearrangements induced by mechanical stimulation have been described during axonal outgrowth (32), and that intracellular Ca^2+^ increments have been shown to promote actin assembly to improve melanoma cell migration (48), we postulated that in nascent adhesion TRPV2 mechanically-induced signaling contributes to F-actin bundles structure stabilization, by promoting cofilin-1 inactivation. Interestingly, in advanced melanoma models, TRPV2 also associates with the intermediate filament vimentin network, conceivably in order to extensively regulate cytoskeletal organization and adhesion structures mechanical maturation (27, 28). Note that as such, vimentin expression often correlates with tumor aggressiveness (49).

As the cell leading edge advances, a subpopulation of nascent adhesions disassembles, and the remainder grow and mature into focal complexes and then FAs. Nascent adhesion growth is accompanied by the recruitment of the mechanosensitive adhesion protein talin, which can bind directly to paxillin but also to actin and integrins (30). In metastatic melanoma models, we observed that TRPV2-mediated Ca^2+^ influx directly regulates calpain activation and the ensuing cleavage of talin. With the Ca^2+^-dependent protease calpain being a major regulator of adhesion components degradation and its substrate, talin, directly impacting the recruitment of the cytoskeletal adapters and the mechanical engagement of FAs (29, 31). To date, the plasma membrane elements controlling the calpains system have been poorly described, raising the question of its mechanosensitive regulation. In metastatic melanoma cells, TRPV2 mechanosensitive channel mediates a Ca^2+^ signal directly activating calpain-mediated proteolysis of talin, hindering adhesion mechanical maturation and favoring its turnover.

In conclusion, CMM, which is traditionally viewed as one of the most metastatic and therapy-resistant malignancy, remains an incurable disease for the great majority of patients and, consequently, is in great need for specifying the molecular mechanisms underpinning metastatic dissemination. Over the last decade, it appeared evident that Ca^2+^ channels act as important regulators of specific steps in tumor progression (4, 7). We hereby reported a central role for the prominently expressed Ca^2+^-conducting TRPV2 channel, during the dynamic process of melanoma cells metastatic dissemination and defined the following model (Figure 7G). By being recruited at the PM within paxillin-rich proximal nascent adhesion structures, TRPV2 mechanosensitive channel lies at the interface between the metastatic cell intracellular machinery and the TME. In highly invasive metastatic melanoma cells, force-induced mechanical signals trigger TRPV2-mediated Ca^2+^ influx, which induces the Ca^2+^-dependent cleavage of talin by the calpains, spatiotemporally regulating cell adhesion dynamics. Concomitantly, TRPV2 enables F-actin stabilization by directly controlling cofilin-1 activity. As a central component of the F-actin/adhesion/ECM interface, mechanically activated TRPV2 coordinates dynamic cytoskeletal rearrangements intertwined with active adhesion site turnover. TRPV2 therefore represents a great molecular candidate for mediating a tunable force-transmitting structural linkage from the cytoskeleton to the TME *via* adhesion complexes, controlling *in fine* melanoma cell migration.

By being directly correlated to the aggressiveness of the tumor and to patient mortality in human melanoma biopsies, TRPV2 stands out as a valuable biomarker for malignant tumors with bad prognosis. Finally, Ca^2+^ channels represent a propitious class of drug targets due to their accessibility to pharmacological modulation and their exposure at the cell surface. TRPV2 pharmacological blockade therefore hints as a promising therapeutic option for migrastatics in the treatment of advanced-stage melanoma.

## Additional information

- This work was funded by Roche/Groupe Cancérologie Cutanée de la Société Française de Dermatologie, the University of Rennes-1, Fondation ARC, La Ligue contre le cancer (Comités 35 & 86), the Région Nouvelle Aquitaine (Chaire Universitaire Canaux Calciques et Mélanome, A.P) and the French Government (ANR, program #ANR-12-JSV2-0004-001). K.F.S was an ANR postdoctoral fellowship recipient. E.B was supported by the Région Poitou-Charente.
- Corresponding author: Aubin PENNA - Université de Poitiers, CNRS, 4CS, PBS, B36, 1 rue G. Bonnet, F-86000 Poitiers, France. +(33) 549 453 996 -
- The authors declare no potential conflicts of interest.
- 5723 words, and a total of 7 figures and tables.

## Acknowledgments

We are grateful to BIOalternatives (FX Bernard) for providing normal human epidermal melanocytes, Dr P-Y Rescan (LPGP-INRA, Rennes, France) for providing access to the zebrafish husbandry and technical guidance, the IRSET Inserm U1085 SMS and DR@TE teams for housing and facility access, as well as to the Abbelight company (N. Bourg) and both the MRic and ImageUP microscopy platforms for critical assistance. We thank the TCGA Research Network. This work would not have been possible without Dr L Larue, Dr MD Galibert and Dr O Destaing help and critical discussions, and without Dr B Jégou’s precious support.

## Author Contributions

Conceptualization, A.P; Formal analysis, K.F.S, E.B, S.L-P; Investigation, K.F.S, E.B, D.LD, A.M, S.M-L, F.R, A.J, R.V and R.P; Methodology, A.P and K.F.S; Project Administration: A.P; Resources, S.T-D; Supervision: S.M-L, A.F, A.K, B.C, S.T-D and A.P; Visualization: K.F.S; E.B; S.L-P and A.P; Writing – Original Draft, K.F.S and A.P; Writing – Review & Editing, K.F.S, S.L-P, S.T-D and A.P; Funding Acquisition, A.P. All authors have read and acknowledged the content of the manuscript.

