## Supplementary methods, tables and figures for "The mechanosensitive TRPV2 calcium channel controls human melanoma invasiveness and metastatic potential"

### Supplemental Information (SI)

#### Supplemental Methods

##### *Additional Bioinformatics analyses*

Gene-centric RMA-normalized TRPV2 mRNA expression data in the Broad Institute and Novartis's Cancer Cell Line Encyclopedia (CCLE) larger cell panel was obtained through the CCLE website (https://portals.broadinstitute.org/ccle) (1). To establish the differential plot of TRPV2 expression across all cancers plus the specific analysis in SKCM compared to healthy tissue we analyzed RNA sequencing expression data of tumors and normal samples from the TCGA and the GTEx projects through the GEPIA web server (http://gepia.cancer-pku.cn/index.html) (2).

##### *Additional Biochemical Techniques*

For Immunoblotting, cells were lysed in RIPA buffer (150 mM NaCl, 50 mM Tris, 0.1% v/v SDS, 0.25% v/v Na-Deoxycholate, 1% v/v NP-40, 1 mM EDTA; pH7.4) supplemented with a protease and phosphatase inhibitor cocktail for 20 min on ice. The protein concentration in supernatant was estimated using the BCA assay according to Pierce's protocol. A total of 30 µg of protein was loaded into NuPage 4-12% gels and transferred onto a nitrocellulose membrane.

Co-immunoprecipitation experiments were performed using cells cultured in 100 mm fibronectin-coated dishes. After sample preparation 1.5 mg of cell lysate per sample was used for immunoprecipitation as described elsewhere (3). Briefly, supernatants were tumbled for 20 min on ice with the appropriate primary antibody, followed by 3h of incubation at 4°C with protein A-Sepharose beads (Sigma Aldrich). The beads were then washed five times with a lysis buffer, suspended in Laemmli sample buffer and boiled for 5 min. The samples were resolved and blotted according to the above-described protocol.

For cell surface proteins biotinylation assays, cells were cultured to 75% confluency in 6-well plate dishes, washed in cold PBS and incubated with the biotinylation reagent (Sulfo-NHS-LC-biotin 0.5 mg/mL in PBS pH8 with 0.1 mM $CaCl_2$, 1 mM $MgCl_2$) for 30 min at 37°C. Free biotinylation reagent was then removed by washing twice in PBS containing 50 mM glycine and 0.1% BSA and once in PBS alone. Cell lysates were prepared as described above in RIPA buffer supplemented with 25 mM $NH_4Cl$. 100 µg of total protein were incubated with 25 µL of High-capacity Streptavidin agarose resin for 1h at 4°C. Captured proteins were eluted in NuPAGE-LDS sample buffer, 4% SDS, 160 mM DTT by heating 10 min at 70°C.

##### *Single cell $Ca^{2+}$ Imaging*

Cells plated on glass coverslips were loaded with 5 µM Fura-2-AM in DMEM at 37°C for 45 min and then washed three times in HBSS solution (142.6 mM NaCl, 5.6 mM KCl, 2 mM $CaCl_2$, 1 mM $MgCl_2$, 10 mM HEPES, 5 mM Glucose, pH 7.4) followed by de-esterification at 37°C for 15 min. Changes in $[Ca^{2+}]_i$ were monitored in cells bath in HBSS using a DMIRB (Leica) inverted microscope-based imaging system equipped with a 40x /1.35 UApo N340 high UV light transmittance oil immersion objective (Olympus), a CoolSnapHQ fast-cooled monochromatic digital camera (Princeton instrument), a DG-4

Ultra High Speed Wavelength Switcher (Sutter Instruments) for fluorophore excitation and METAFLUOR software (Universal Imaging) for image acquisition and analysis. Data were acquired every 10s (emission at 510 nm) at 340 and 380 nm excitation wavelengths and all images were background-subtracted. Cells with spontaneous or aberrant $Ca^{2+}$ activity were identified by imaging and eliminated from the analysis. Depicted curves represent a minimum of 3 independent experiments.

***Immunofluorescence staining and confocal microscopy.***

Glass coverslips were coated with human fibronectin (50 µg/mL) in $Ca^{2+}$/$Mg^{2+}$-PBS. WM266.4 and 451Lu cells seeded at low confluency were fixed in PBS-4% PFA-4% sucrose for 10 min, quenched with PBS-glycine 0.1 M for 30 min, then permeabilized and blocked with PBS-10% FCS-0.2% saponin for 15 min. Primary antibodies diluted in PBS-5% FCS-0.2% saponin incubation lasted 2h. For immunodetection, secondary antibodies (diluted 1/1000) were incubated for 30 min at 37°C. F-actin was detected using Acti-stain. Nuclei were stained with DAPI at 1 µg/mL for 15 min. Coverslips were mounted in Vectashield. Fluorescence micrographs were taken using laser scanning confocal systems (TCS SP8 model mounted on a DMI 6000 CS inverted microscope, Leica or IX81-based Olympus FV1000). For FAs counting (vinculin), fixation was done in 70% ethanol and no saponin was used.

***Single molecule imaging***

**Optical setup.** We used an Olympus IX83 inverted microscope with an autofocus system. The excitation path was composed of three laser lines: 637 nm, 532 nm and 405 nm (Errol lasers) and a TIRF module (Errol lasers) used in combination with a matched 390/482/532/640 multiband filter (LF405/488/532/635-A-000, Semrock). The fluorescence was collected through an Olympus x100 1.49 NA oil immersion objective lens. The detection path was composed of a SAFe module (Abbelight) and a Flash 4 v3 (Hamamatsu). The pixel size in the object was 100 nm.

**Image acquisition.** The diffraction limited epifluorescence images were acquired at low illumination irradiance (0.15 $kW.cm^{-2}$), while the dSTORM images were obtained using a high illumination irradiance (4 $kW.cm^{-2}$) until a sufficient molecule density was obtained (around 1 molecule per $µm^2$) and the acquisition could be started. The exposure time was set at 50 ms. Acquisitions were performed using the Nemo software (Abbelight). To achieve a single molecule regime in dSTORM acquisition, a dedicated buffer (Smart kit, Abbelight) was used.

**Image processing.** Acquired data were processed using the Nemo software (Abbelight). After removing the background signal, molecules were detected and the numbers of EPI and UAF photons were measured to extract the corresponding axial positions. Lateral drifts were corrected from the localized data thanks to a cross-correlation based algorithm.

***Cell Proliferation***

Cell viability was evaluated using the MTT assay. Briefly, 40,000 cells were cultured for 24h in flat-bottom 96-well plates in a final volume of 100 µL. Then, 15 µL of MTT (5 mg/mL in PBS) solution were added and after 4h of incubation at 37°C the absorbance was measured at 570 nm using the EnSpire® 2300 Multilabel Plate Reader (Perkin Elmer). Proliferation was also assessed by cell counting: briefly, the cells were cultured in 6-well dishes in duplicate at a density of 5,000 or 20,000 cells per well in 2 mL of medium supplemented with or without FBS. The medium was changed every 48h. Cell number at the indicated time points was determined by counting using a hemocytometer.

***Three-dimensional (3D) Spheroid Invasion Assay***

Melanoma spheroids were prepared using the liquid overlay method. Briefly, 24-well culture plates (Corning) were coated with 1.5% agarose (Life Technologies) in sterile water. Cells from a single-cell suspension were added at 10,000 per well. The plates were incubated at 37°C in a 5% CO2 atmosphere until spheroids were formed (72h). Spheroids were harvested and implanted into a gel of rat collagen I (Becton Dickinson). Complete melanoma medium was overlaid on top of the solidified collagen. Pictures of the invading spheroids were taken each day using an inverted microscope. Images were processed and growth areas measured using ImageJ software.

***Calpain activity.***

Calpain activity was measured using the calpain activity assay kit (Promokine). Briefly $2 \times 10^6$ cells were seeded on 6-well plates for 12h, washed and suspended in 100 µL of extraction buffer. 200 µg of proteins were diluted in 85 µL of extraction buffer. Calpain activity was revealed by adding 10 µL of reaction buffer and 5 µL of fluorescent calpain substrate. Activity was measured using a fluorometer equipped with a 400 nm excitation filter and a 505 nm emission filter. 1 µL of calpain inhibitor was added to subtract the background.

***Zebrafish tumor cell implantation and micrometastasis analysis***

Zebrafish embryos were raised, staged and maintained according to standard procedure. For cell microinjection, 2 days-post fertilization (dpf), phenylthiourea (PTU, Sigma-Aldrich)-treated zebrafish were dechorionized and anesthetized using 0.04 mg/mL Tricaine. WM266.4 human melanoma cells harvested in HyQTase™, counted and labelled or not with CM-Dil tracker were then mixed in equal quantity and loaded into borosilicate capillaries at a density of $8 \times 10^7$ cells/mL. Injections were performed using a pneumatic picopump (World Precision Instruments) and a micromanipulator. Cell injection was performed above the ventral duct of Cuvier as described in (4). After confirmation of a visible cell mass at the injection site, selected zebrafish were transferred to an incubator and maintained at 34°C for 36h. Micrometastasis formation was analyzed on living zebrafish embryos anesthetized with Tricaine. Images were captured with a DeltaVision imaging system and processed using ImageJ/Fiji software.

***Immunostaining of metastasis from xenografts experiments***

For metastases immunostaining, lungs were perfused with PBS, fixed in 4% paraformaldehyde, paraffin-embedded and sectioned. Sections of lungs (4 µm) were treated on a Discovery Automated IHC stainer (Roche) to remove paraffin, unmask epitopes and sequentially stain with rabbit polyclonal antibodies against TRPV2 (1:50; HPA044993 Sigma) or pmel17/HMB45 (1:50) and rat monoclonal antibody anti-mouse CD31 (1:200; clone SZ31) before detection with Alexa Fluor-conjugated secondary antibodies. Sections were mounted in Antifade Reagent with DAPI (Life Technologies).

#### Supplemental tables

##### *List of primers*

Each primer pair was designed and validated *in silico* using primer3plus and primer-blast software (http://www.bioinformatics.nl/cgi-bin/primer3plus/primer3plus.cgi, http://www.ncbi.nlm.nih.gov./tools/primer-blast/). Primers were designed to localize at intron/exon junctions of sequences commune to all variants of each gene.

| Gene | Amplicon size | Forward primer sequence (5'-3') | Reverse primer sequence (5'-3') |
|---|---|---|---|
| TBP | 161 | CGGAGAGTTCTGGGATTGT | GGTTCGTGGCTCTCTTATC |
| TRPV1 | 120 | GACCTGTGCCGTTTCA | CCTGTGCGACGTGGACTCA |
| TRPV2 | 199 | GGAATACACAGAGGGCTCCA | CCTCTTCTCAATGGCGATGT |
| TRPV3 | 226 | ACGAGGCAACAACATCCTTC | CCGCTTCTCCTTGATCTCAC |
| TRPV4 | 190/370 | CCCGTGAGAACACCAAGTTT | AGTTCATTGATGGGCTCCAC |
| TRPV6 | 208 | GGATCTGCGGACGGGAGTA | CGAGACACTGAGGGCATAGGA |
| TRPC1 | 201 | TGGGATGATTTGGTCAGACA | TCTGCCACCAGTGTAGGATG |
| TRPC6 | 121 | CCTTGCTGTTGCCATTGGAC | GAAGGAGGCTGCGTGTGCTA |
| TRPM2 | 303 | GGCAGTGGAAGCCTTCAGAT | GATAAAGCGGCTGCGTGAAG |
| TRPM7 | 226 | AATAATCGGAGGTCTGGCCG | AGCGCTTGGTTTCTGGATCA |
| TRPA1 | 140 | TGCATGTTGCATTCCACAGAAG | TTGAGGGCTGTAAGCGGTTCATA |
| STIM1 | 109 | TGTGGAGCTGCCTCAGTATG | CTTCAGCACAGTCCCTGTCA |
| STIM2 | 114 | GTCTCCATTCCACCCTATCC | GGCTAATGATCCAGGAGGTT |
| ORAI1 | 161 | ATGAGCCTCAACGAGCACT | GTGGGTAGTCGTGGTCAG |
| SK3 | 174 | TGGACACTCAGCTCACCAAG | GTTCCATCTTGACGCTCCTC |
| BRN2 | 98 | GTAACTGTCAAATGCGCGGC | GAGGTGAGCAGGCTGTAGTG |

Note that the TRPV4 specific primer pair use for RT-PCR amplify TRPV4 variants 1, 4 and 5 with an amplicon size of 370 bp and TRPV4 variants 2 and 3 with an amplicon size of 190 bp (5). For qPCR experiments, an alternative primer pair was used amplifying all variants with the same amplicon size of 76 bp (Fwd 5'-CTACGCTTCAGCCCTGGTCTC-3'; Rev 5'-GCAGTTGGTCTGGTCCTCATTG-3' (6)).

***Antibodies & Reagents***

| ---------- *Antibodies* ------------ |  |  |  |
|---|---|---|---|
| ß-actin | mouse monoclonal | clone AC-74 | Sigma |
| Talin | mouse monoclonal | clone 8d4 | Sigma |
| Vinculin | mouse monoclonal | clone VLN01 | ThermoFisher |
| PMEL (pmel17/HMB45) | rabbit polyclonal | | ThermoFisher |
| TRPV2 | rabbit polyclonal | VRL-1 H-105 | Santa Cruz Biotechnology |
| Paxillin | mouse monoclonal | clone 349 | BD |
| Phospho-FAK Tyr397 | rabbit polyclonal | | Cell Signaling |
| Phospho-p44/42 MAPK (Erk1/2) (Thr202/Tyr204) | rabbit monoclonal | clone D13.14.4[E] | Cell Signaling |
| Cofilin<br>Phospho-Cofilin Ser3 | rabbit monoclonal | clones D3F9<br>77G2 | Cell Signaling |
| Vimentin | mouse monoclonal | clone V9 | DAKO |
| CD31/PECAM-1 | rat monoclonal | clone SZ31 | Dianova |
| Goat-anti-rabbit Alexa Fluor-488 or -555<br>Goat anti-mouse Alexa Fluor-488 or -555 | | | Invitrogen |

| ---------- *Reagents* ------------ |  |
|---|---|
| SIR-actin | Spirochrome |
| Acti-stain 555 Phalloidin | Cytoskeleton |
| Rhodamine Phalloidin | ThermoFisher |
| Vectashield | Vectorlabs |
| Fura-2/AM | Molecular Probes |
| (-)-Cannabidiol | Tocris |
| NuPage 4-12% gels | ThermoFisher |
| Nitrocellulose membrane | GE Healthcare |
| Bicinchoninic acid (BCA) protein assay kit<br>Sulfo-NHS-LC-Biotin | Pierce |
| High Capacity Streptavidin Agarose Resin | Invitrogen |
| ECL RevelBot | Ozyme |
| human fibronectin<br>protease and phosphatase inhibitor cocktail | Sigma |
| All other chemicals | Sigma-Aldrich |

| *Cell lines & Culture reagents.* | | |
|---|---|---|
| | | **Growth medium** |
| Human melanocytes | BIOalternative, | a |
| 501mel cells | S. Tartare-Deckert, | b |
| WM115<br>WM266.4 | ATCC , | c |
| WM164<br>451Lu<br>WM2032 | WISTAR institute, | b<br>b<br>a |
| NCI-60 collection | Charles River Laboratories, | b |

a. 80% MCDB153 medium + 20% Leibovitz's L-15 medium + 1.68 mM $CaCl_2$ + 2% hi FCS.
b. DMEM medium + 8% hiFCS + 2 mM L-Glutamine
c. RPMI1640 + glutamax + 8% hiFCS

| *37°C - 5% $CO_2$ atmosphere* | |
|---|---|
| All Media<br>L-glutamine<br>Glutamax<br>0.25% trypsin-EDTA | Gibco |
| Fetal calf serum<br>heat-inactivated (hiFCS) | Eurobio |
| DMSO (10% in growth medium, for cryopreservation) | Sigma |
| MycoAlert detection kit | Lonza |

#### Supplemental Figures with Legends

**Supplemental Figure 1 (relative to Figure 1). TRPV2 is predominantly expressed in melanoma cells compared to cancer cells from other tissue origins**

(A) TRPV2 mRNA expression analysis within the Broad-Novartis Cancer Cell Line Encyclopedia (CCLE) project RNAseq data. The box plot is sorted and colored by average distribution of TRPV2 expression in a lineage. Lineages are composed of cell lines from the same area/organ or system of the body. The number of cell lines per lineage is indicated. The highest average distribution is in red on the left. The dashed line within a box is the mean.

(B) GEPIA (Gene expression profiling interactive analysis) generated dot plot profiling TRPV2 gene expression across multiple cancer types and paired normal samples (TCGA tumors *vs* TCGA normal) based on TCGA RNAseq data directly extracted from tissue samples. Cancer types are as follow (T, tumor sample; N, matched normal sample): ACC, Adrenocortical carcinoma (T n=77; N n=-); BLCA, Bladder carcinoma (T n=404; N n=19); BRCA, Breast invasive carcinoma (T n=1085; N n=112); CESC, Cervical squamous cell carcinoma and endocervical adenocarcinoma (T n=306; N n=3); CHOL, Cholangiocarcinoma (T n=36; N n=9); COAD, Colon adenocarcinoma (T n=275; N n=41); DLBC, Diffuse Large B-cell Lymphoma (T n=47; N n=-); ESCA, Esophageal carcinoma (T n=182; N n=13); GBM, Glioblastoma multiform (T n=163; N n=-); HNSC, Head and neck squamous cell carcinoma (T n=519; N n=44); KICH, Kidney chromophobe (T n=66; N n=25); KIRC, Kidney renal clear cell carcinoma (T n=523; N n=72); KIRP, Kidney renal papillary cell carcinoma (T n=286; N n=32); LAML, Acute Myeloid Leukemia (T n=173; N n=-); LGG, Lower grade glioma (T n=518; N n=207); LIHC, Liver hepatocellular carcinoma (T n=369; N n=50); LUAD, Lung adenocarcinoma (T n=483; N n=59); LUSC, Lung squamous cell carcinoma (T n=486; N n=50); MESO, Mesothelioma (T n=87; N n=-); OV, Ovarian serous cystadenocarcinoma (T n=426; N n=-); PAAD, Pancreatic adenocarcinoma (T n=179; N n=4); PCPG, Pheochromocytoma and Paraganglioma (T n=182; N n=3); PRAD, Prostate adenocarcinoma (T n=492; N n=52); READ, Rectum adenocarcinoma (T n=92; N n=10); SARC, Sarcoma (T n=262; N n=2); SKCM, Skin cutaneous melanoma (T n=461; N n=1); STAD, Stomach adenocarcinoma (T n=408; N n=36); TGCT, Testicular germ cell tumor (T n=137; N n=-); THCA, Thyroid carcinoma (T n=512; N n=59); THYM, Thymoma (T n=118; N n=2); UCEC, Uterine corpus endometrial carcinoma (T n=174; N n=13); UCS, Uterine carcinosarcoma (T n=57; N n=-); UVM; Uveal melanoma (T n=79; N n=-).

A **Broad-Novartis Cancer Cell Line Encyclopedia (CCLE) project RNAseq data**

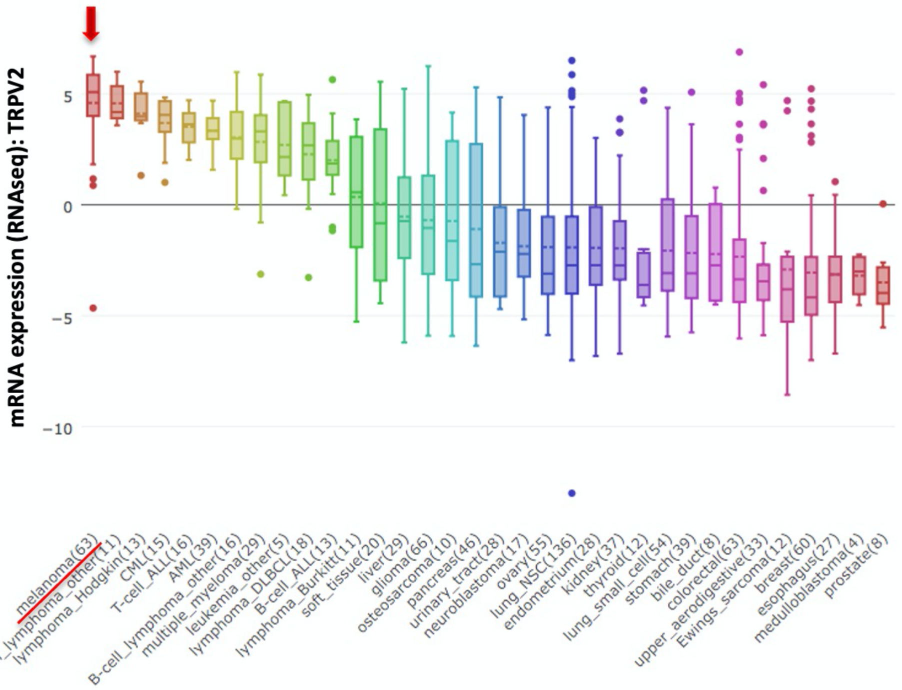


B **The Cancer Genome Atlas (TCGA) cohort RNAseq data.**

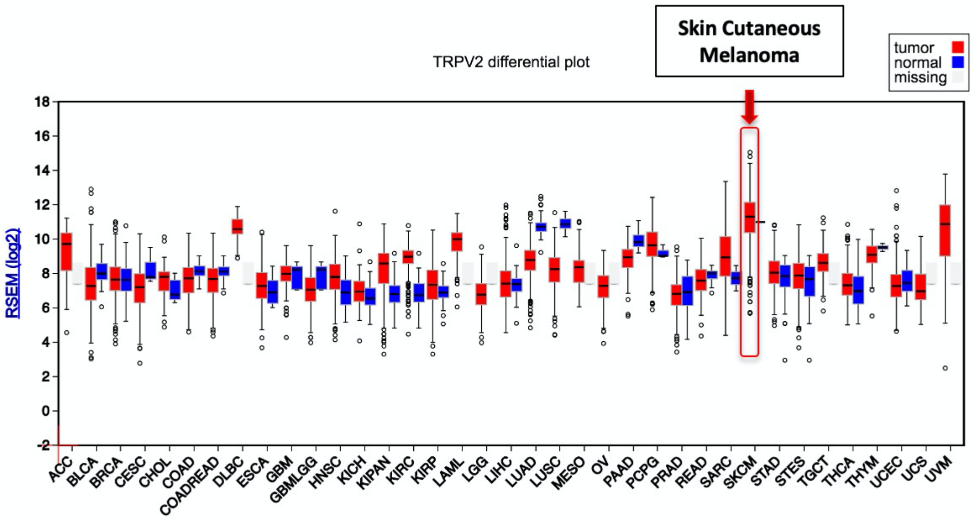

**Supplemental Figure 2 (relative to Figure 2). TRPV2 expression, plasma membrane localization and contribution to the basal $Ca^{2+}$ influx correlate with the invasive phenotype of melanoma cells**

(A) Representatives $Ca^{2+}$ influx responses to 40 µM Cannabidiol (CBD) stimulation. Data represents means ± SD from at least 20 cells.

(B) Confocal microphotographs showing the distribution of TRPV2 (red) and stained nuclei (blue) in the indicated melanoma cell lines. The dotted line highlights the membrane edge in the WM793 cell line. Note that TRPV2 distribution at the plasma membrane correlates with the invasive potential of the melanoma cell lines.

(C) Representative kinetics of CBD induced $Ca^{2+}$ influx in melanoma cell lines modified for TRPV2 expression. Non-invasive 501mel cells (left) overexpressing GFP (Ctl) or GFP-TRPV2. WM266.4 (middle) and 451Lu (right) metastatic cells transduced with either control (Ctrl) or TRPV2-targeting sequences V2-1 and V2-2 shRNAs.

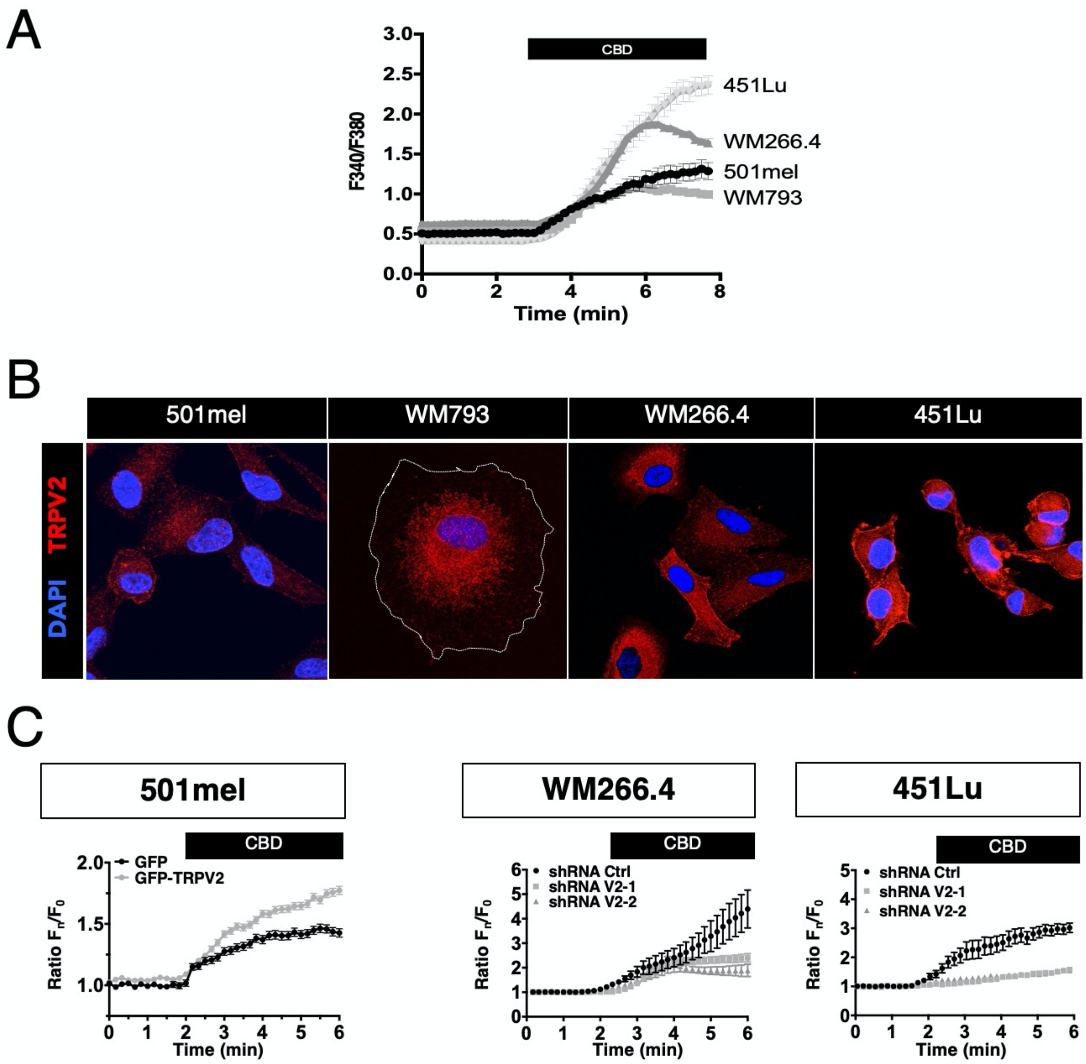

**Supplemental Figure 3. TRPV2 is dispensable for melanoma tumor cells proliferation**

(A) Representative proliferation curves comparing the effect of TRPV2 overexpression in 501mel cells, or repression in WM266.4 and 451Lu cells, measured by MTT at 12, 24, 48 or 72h.

(B) Immunoblotting (IB) of TRPV2, Thr202/Tyr204 Phospho-p44/42 MAPK (pERK) and the β-actin as a loading control in melanoma cell lines modified for TRPV2 expression.

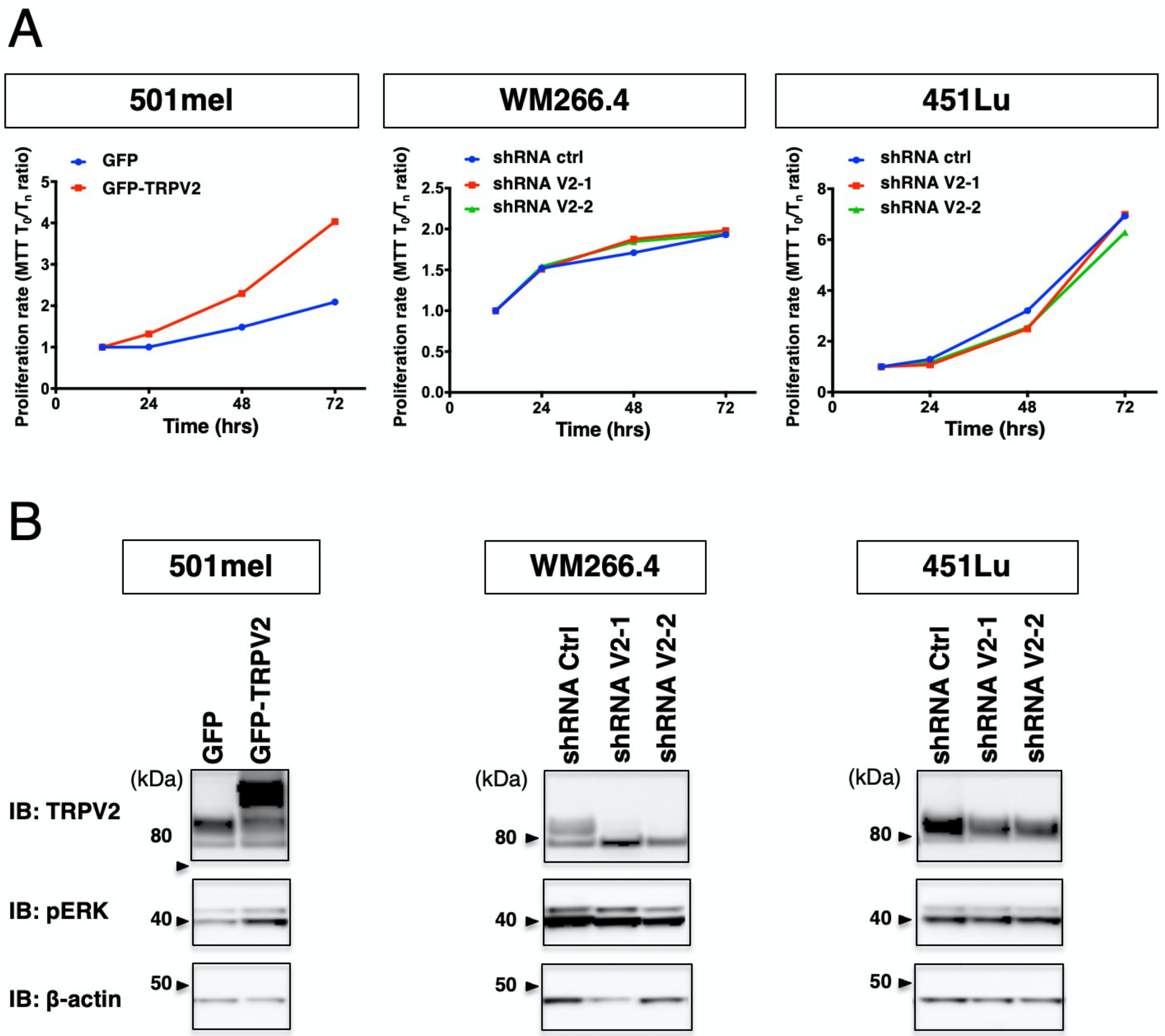

**Supplemental Figure 4 (relative to Figure 3). TRPV2 is essential for melanoma tumor cell migration and invasion**

(A) Schematic representation of 2D cell migration assays using WM266.4 cells transduced with control (red) or TRPV2 (green) shRNAs, and placed in a 0-5% FCS gradient using an Ibidi migration chamber. Representative fluorescent image showing a typical field of seeded cells being evaluated for 12h, under temperature and $CO_2$ controlled conditions.

(B) Invasion kinetics of collagen-embedded spheroids based on Figure 3 D-E experiments.

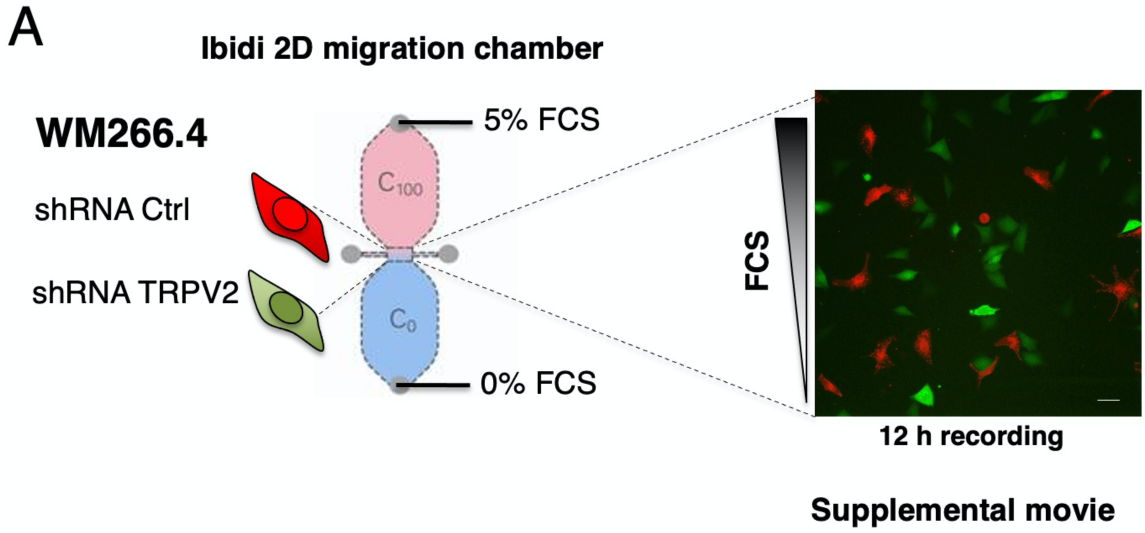


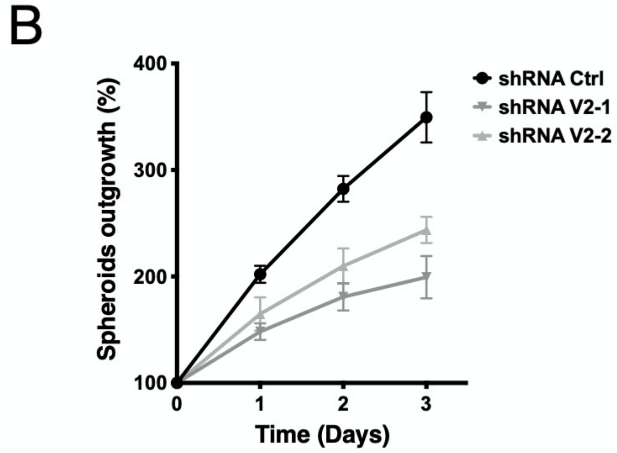

**Supplemental Figure 5 (relative to Figure 4). TRPV2 associates with immature acto-adhesive structures in metastatic cells**

(A) Super-resolution imaging (dSTORM) revealed clustering of TRPV2 channels with paxillin, but not vinculin in migrating WM266.4 cells. a and b insets show expanded view of a region of the cell and arrows highlight TRPV2 channel and paxillin co-clusters.

(B) Representative confocal images of low confluence WM266.4 and 451Lu melanoma cells seeded on fibronectin-coated coverslips. Cell nuclei are depicted in blue (DAPI), TRPV2 in red and vimentin (VIM) in green. Insets show magnification of the indicated area and arrows indicate sites of co-localization.

(C) Representative immunoblotting (IB) of reciprocal immunoprecipitation (IP) of TRPV2 with vimentin in WM266.4 and 451Lu cells. Experiments were performed three times.

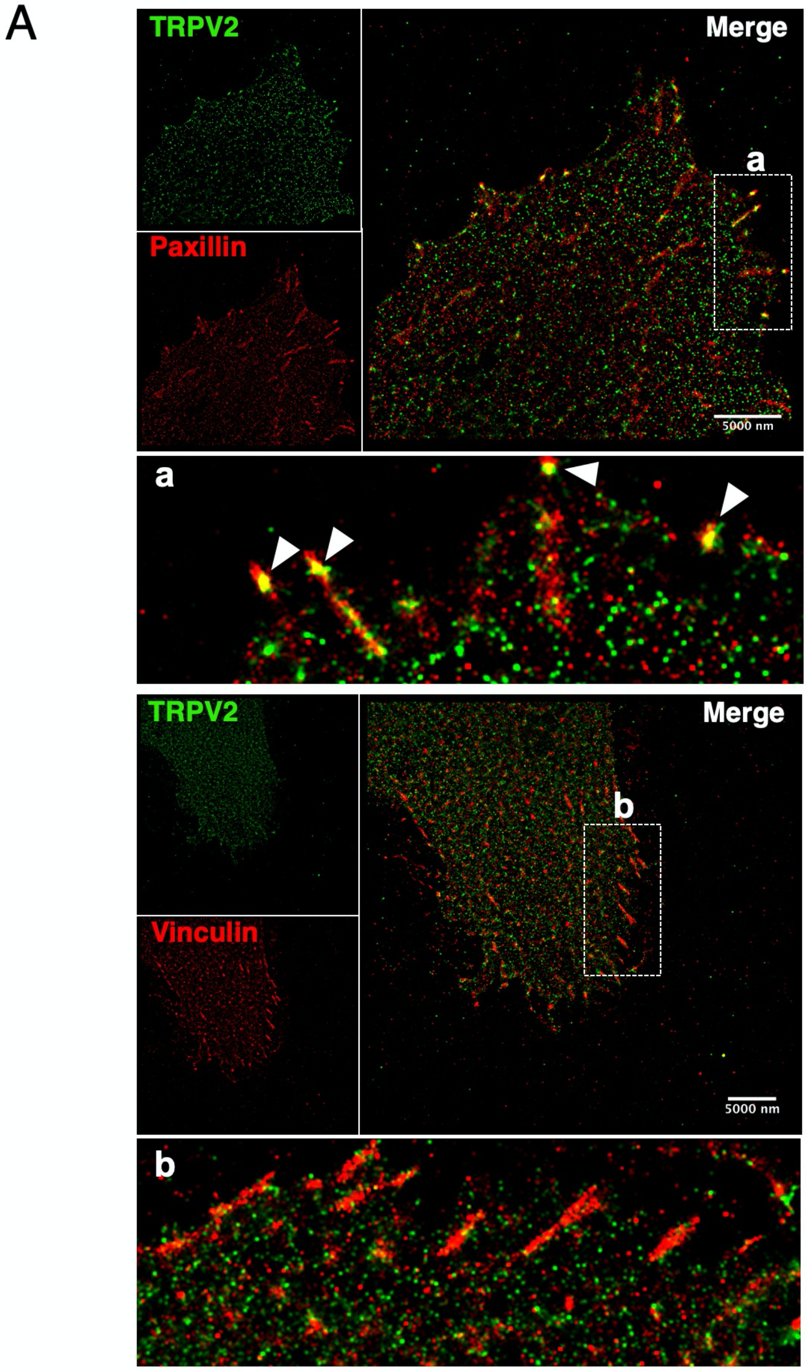

B

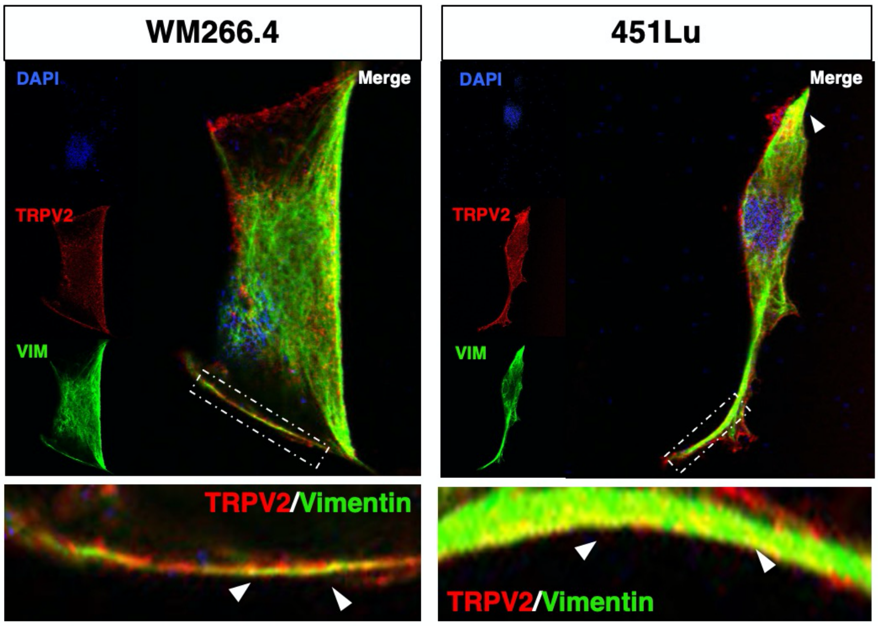
WM266.4
451Lu
DAPI
TRPV2
VIM
Merge
DAPI
TRPV2
VIM
Merge
TRPV2/Vimentin
TRPV2/Vimentin

C

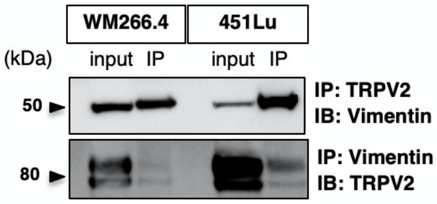
WM266.4
451Lu
(kDa)
input IP
input IP
50
80
IP: TRPV2
IB: Vimentin
IP: Vimentin
IB: TRPV2

**Supplemental Figure 6. Modulating TRPV2 expression does not affect pseudo-EMT markers**

Representative immunoblots analysis of the indicated "epithelial" and "mesenchymal" markers in control (GFP) or GFP-TRPV2 overexpressing 501mel cells, and in WM266.4 and 451Lu cells expressing either a control (ctrl) or a TRPV2-targeting (V2-1 or V2-2) shRNAs. HSP60 was used as a loading control. MITF= Microphthalmia-associated transcription factor; E-CAD= E-Cadherin; FN1= Fibronectin; N-CAD= N-Cadherin; SPARC= secreted protein acidic and rich in cysteine; a-SMA= alpha-smooth muscle actin; VIM= Vimentin.

The star indicates the detection of non-phospho, active β-catenin (β-CAT) (non-phosphorylated on Ser33, Ser37 and Thr41 residues, D13A1 Rabbit mAb, Cell Signalling). Note that 501mel cells display high levels of active β-catenin, acting as a suppressor of cell invasion in melanoma (7). WM266.4 cells strongly express all the mesenchymal markers tested, suggesting that these cells have completed a pseudo-EMT process, while 451lu cells present an intermediate phenotype.

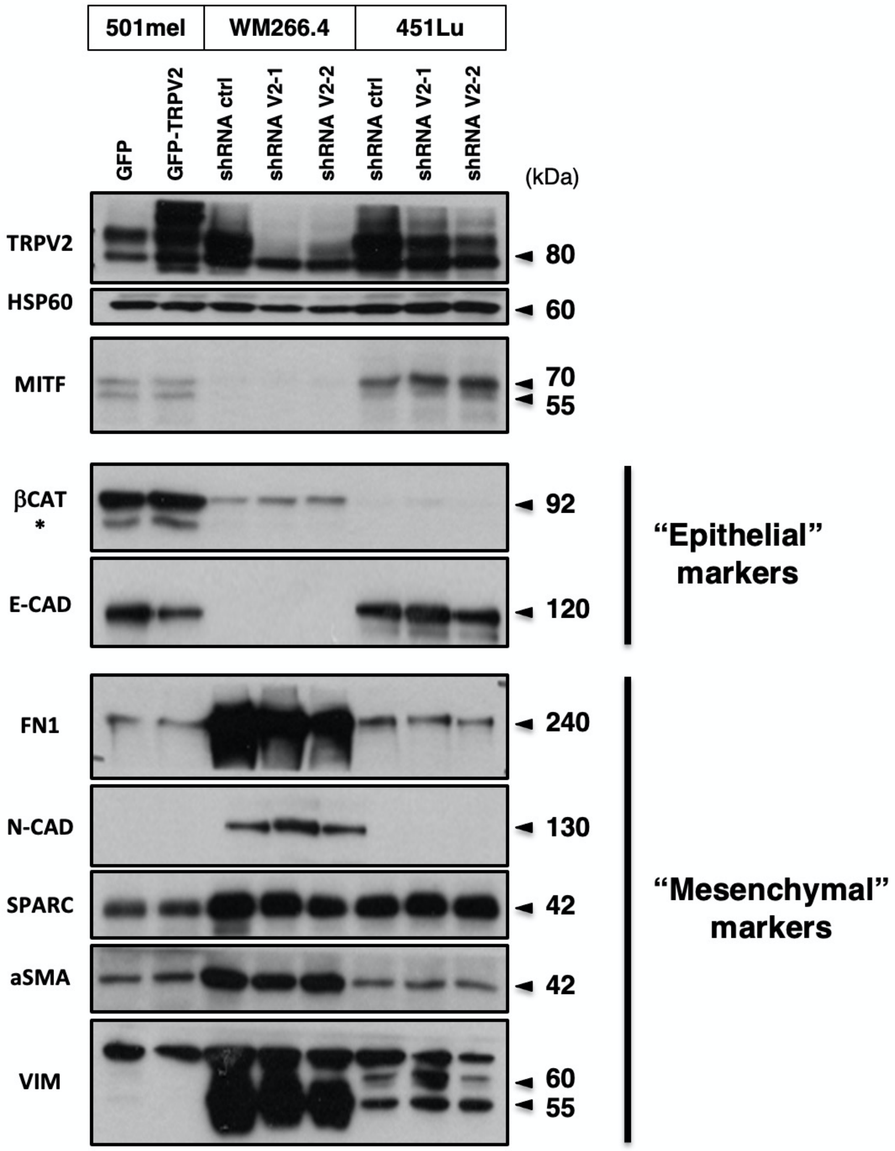

**Supplemental Figure 7 (relative to Figure 5). TRPV2 controls focal adhesions maturation and turnover through the $Ca^{2+}$-dependent activation of calpain**

(A) Representative vinculin (red) and nuclei (DAPI, blue) staining, used to quantify mature FAs per cell, in cell lines where TRPV2 expression was modulated.

(B) Immuno-blotting (IB) assessing the total amount of vinculin following TRPV2 genetic manipulation. β-actin was used as a loading control.

(C & D) IB assessment of the $Ca^{2+}$-dependency of talin cleavage, an adhesion protein proteolyzed by calpain. Transfected 501mel cells (C), or wild-type WM266.4 and 451Lu cells (D) were bathed in nominally $Ca^{2+}$-free (0) or 2 mM $Ca^{2+}$ (2) HBSS external solution. Five minutes prior lysis, cells were treated with either 5 µM Ionomycin, a selective potent $Ca^{2+}$ ionophore allowing normalization of the intracellular [$Ca^{2+}$] with the extracellular [$Ca^{2+}$], or with cannabidiol (CBD, 40 µM for 10 min) eliciting a TRPV2-mediated $Ca^{2+}$ influx. Note that following either of these treatments (both resulting in an increased cytosolic [$Ca^{2+}$]), talin was majoritarly present under its full-length form in 0 mM $Ca^{2+}$ cultured cells, whereas a strong occurrence of the 190 kDa calpain-cleaved isoform was observed in the 2 mM $Ca^{2+}$ external solution bathed cells. In the context of TRPV2-dependent $Ca^{2+}$ influx mediated by CBD stimulation, 501mel cells overexpressing GFP-TRPV2 were used instead of GFP cells lacking TRPV2. In all TRPV2-expressing melanoma cells, CBD-induced $Ca^{2+}$ influx leads to talin proteolysis evoking calpain activation.

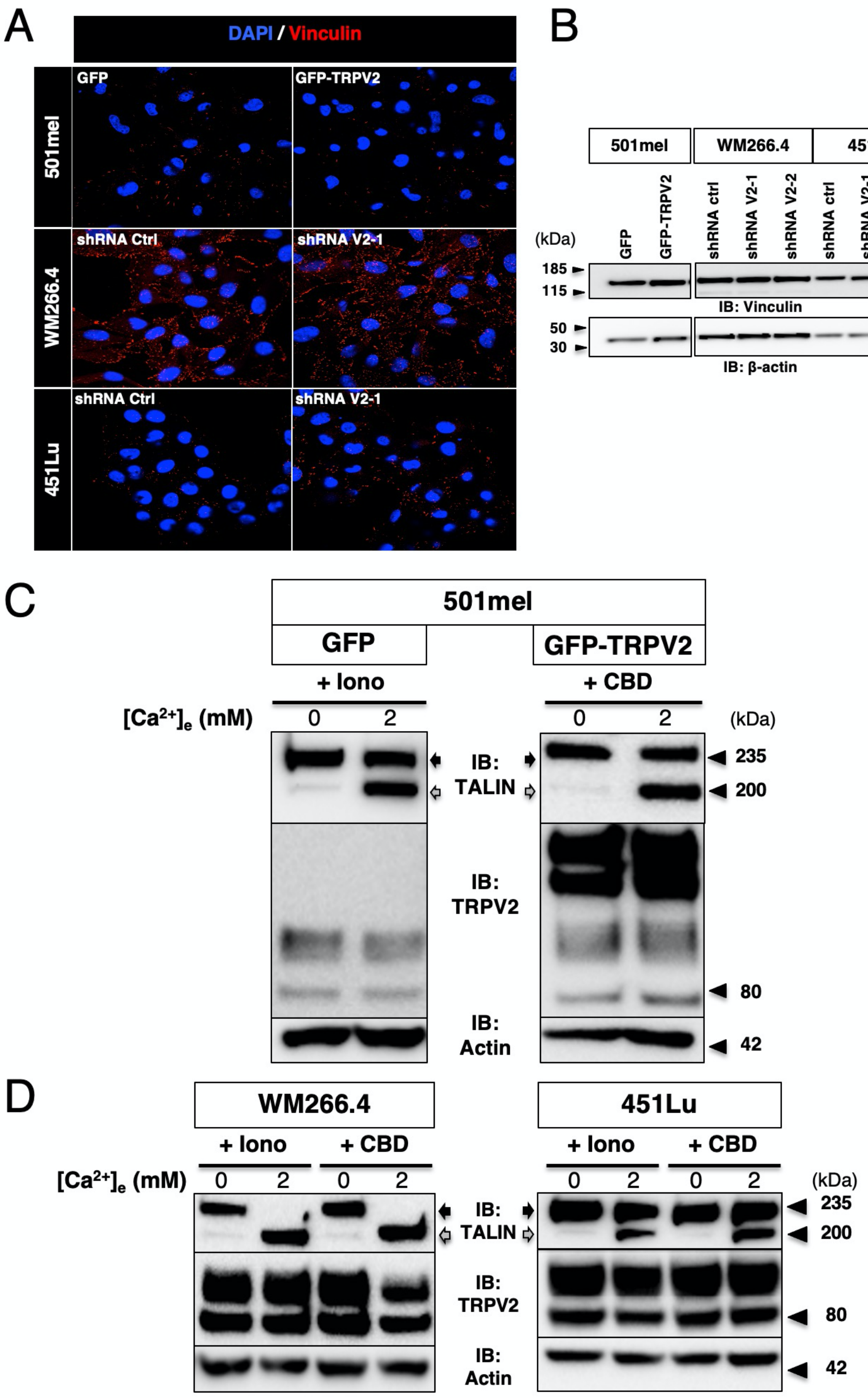

A
DAPI / Vinculin
501mel
GFP
GFP-TRPV2
WM266.4
shRNA Ctrl
shRNA V2-1
451Lu
shRNA Ctrl
shRNA V2-1
B
501mel
WM266.4
451Lu
(kDa)
GFP
GFP-TRPV2
shRNA ctrl
shRNA V2-1
shRNA V2-2
shRNA ctrl
shRNA V2-1
shRNA V2-2
185
115
IB: Vinculin
50
30
IB: β-actin
C
501mel
GFP
GFP-TRPV2
+ Iono
+ CBD
[Ca2+]e (mM)
0
2
0
2
(kDa)
IB: TALIN
235
200
IB: TRPV2
80
IB: Actin
42
D
WM266.4
451Lu
+ Iono
+ CBD
+ Iono
+ CBD
[Ca2+]e (mM)
0
2
0
2
0
2
0
2
(kDa)
IB: TALIN
235
200
IB: TRPV2
80
IB: Actin
42

**Supplemental Figure 8. TRPV2 regulates actin cytoskeleton remodeling and cofilin activation**

(A & B) Fluorescent microscopy images showing live F-actin polymerization in control (GFP) or GFP-TRPV2 overexpressing 501mel cells (A), and in control (Ctrl) or TRPV2-targeting shRNAs transduced WM266.4 cells (B). Quantification of mean fluorescent intensity (top plot) and total cell area (bottom plot) in both populations (*P<0.05, ns=non-significant).

(C) Reverse co-immunoprecipitation (IP) experiments showing a physical interaction between TRPV2 and cofilin in the WM266.4 and 451Lu metastatic melanoma cell lines. Co-IP experiments were performed three times and the immunoblots (IB) show typical results.

(D) Immunoblot assessment of cofilin activation by measuring total and Ser3-phosphorylated cofilin levels in control (shRNA Ctrl) and TRPV2-silenced (shRNA V2) WM266.4 and 451Lu cells.

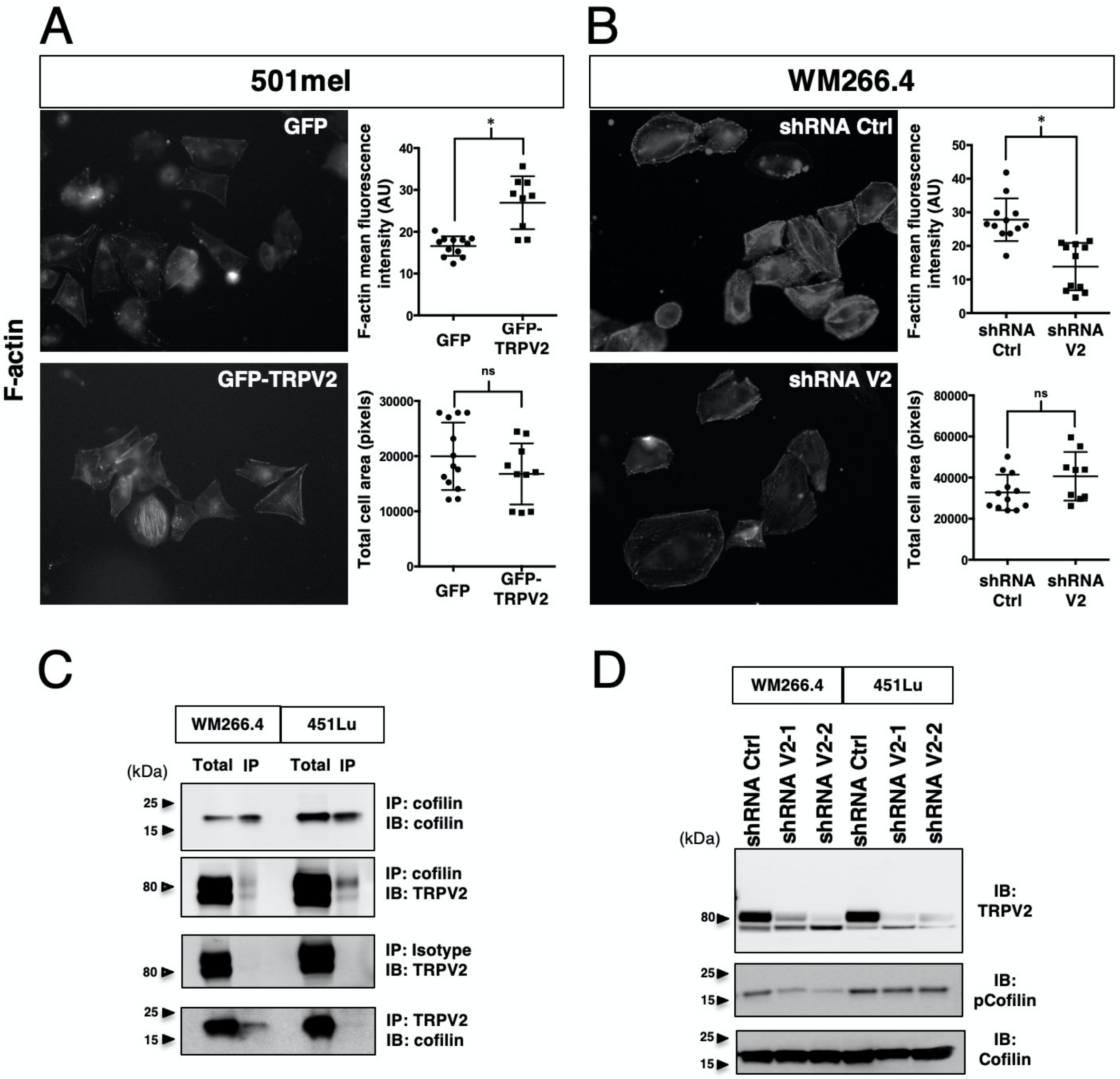

**Supplemental Figure 9 (relative to Figure 6). TRPV2 expression level determines the *in vivo* metastatic potential of melanoma tumor cells**

(A) *Ex vivo* BLI measurements for each necropsy collected organ of representative mice xenografted with either GFP control or GFP-TRPV2 overexpressing 501mel-LUC cells.

(B) Bar graph shows the number of metastatic foci per organ counted at necropsy of mice xenografted with GFP (black) or GFP-TRPV2 (gray) overexpressing 501mel-LUC cells.

(C) Lung extravasation/colonization analysis 24 h after inoculation of 451Lu-LUC cells expressing either ctrl shRNA or TRPV2 targeting shRNA. Scatter plots (Box and whiskers) of normalized BLI photon flux in lungs. Dot, single mouse, n=7 per group.

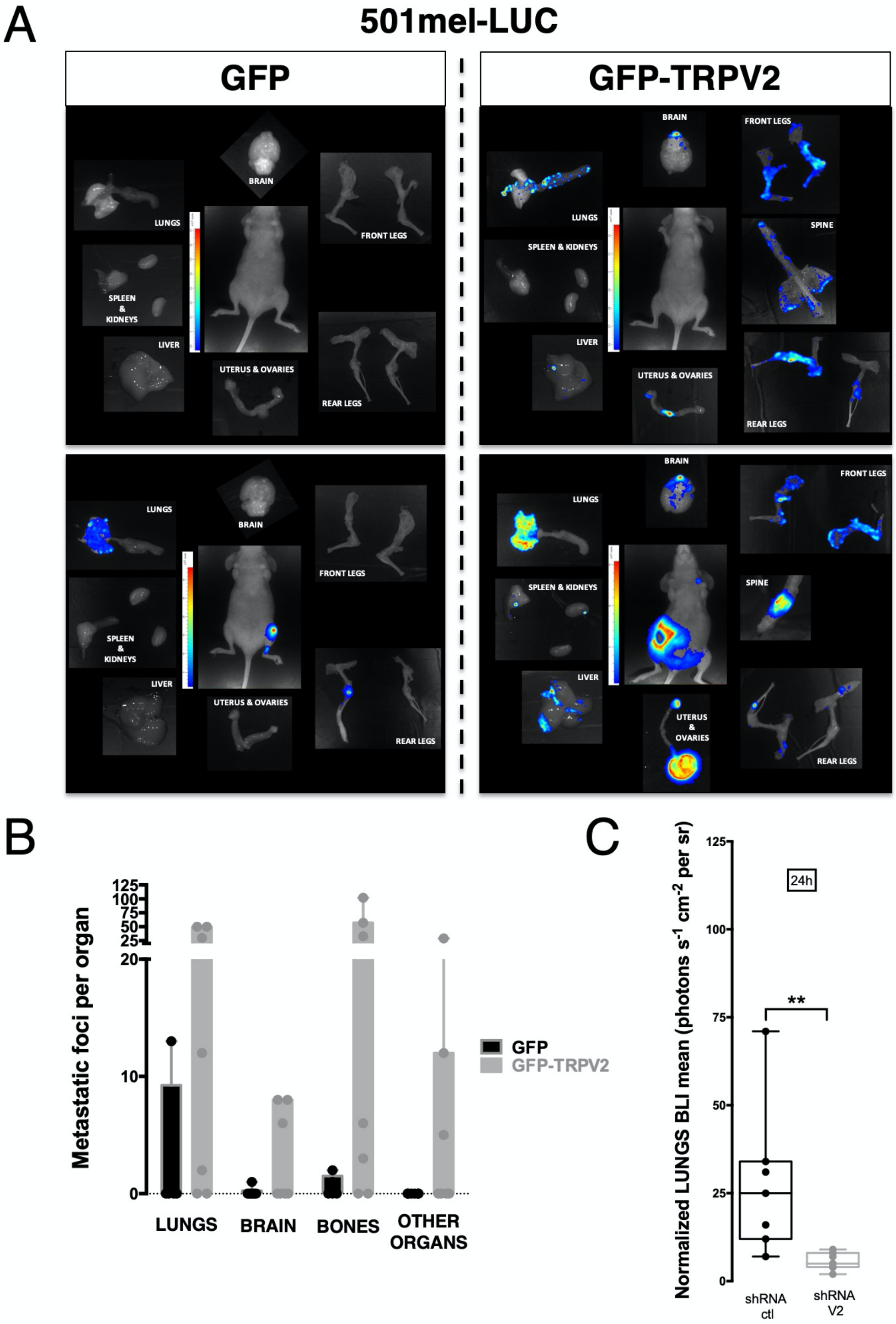

**Supplemental Figure 10 (relative to Figure 6). *Ex vivo* characterization of lung metastasis colonized by melanoma cells**

(A & B) Immunofluorescence analyses of lung sections from mice injected with 451Lu-LUC cells expressing either control or TRPV2 targeting shRNAs. Representative microscopy images of lung sections labelled with DAPI (blue), antibodies against the mouse endothelial cells marker CD31/PECAM-1 (red) and the human melanoma marker MEL (A) or TRPV2 (B) (yellow). Scale bar, 5 mm.

(C) High magnification pictures of lung metastatic foci showing that melanoma cells (a, b insets) have colonized the lung parenchyma, as well as the strong expression and plasma membrane localization of endogenous TRPV2 (c, d insets) in these tumor xenografts.

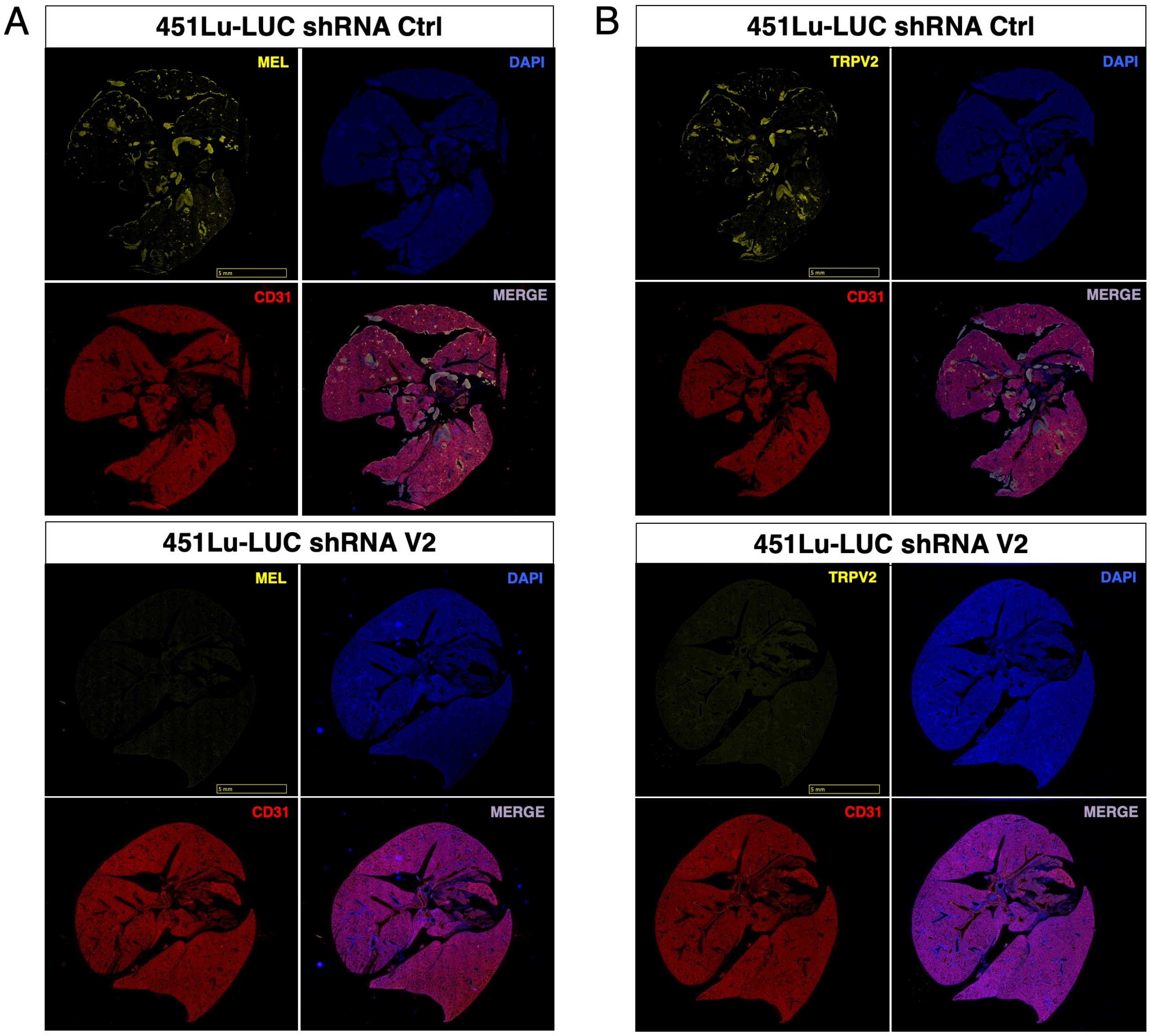

C

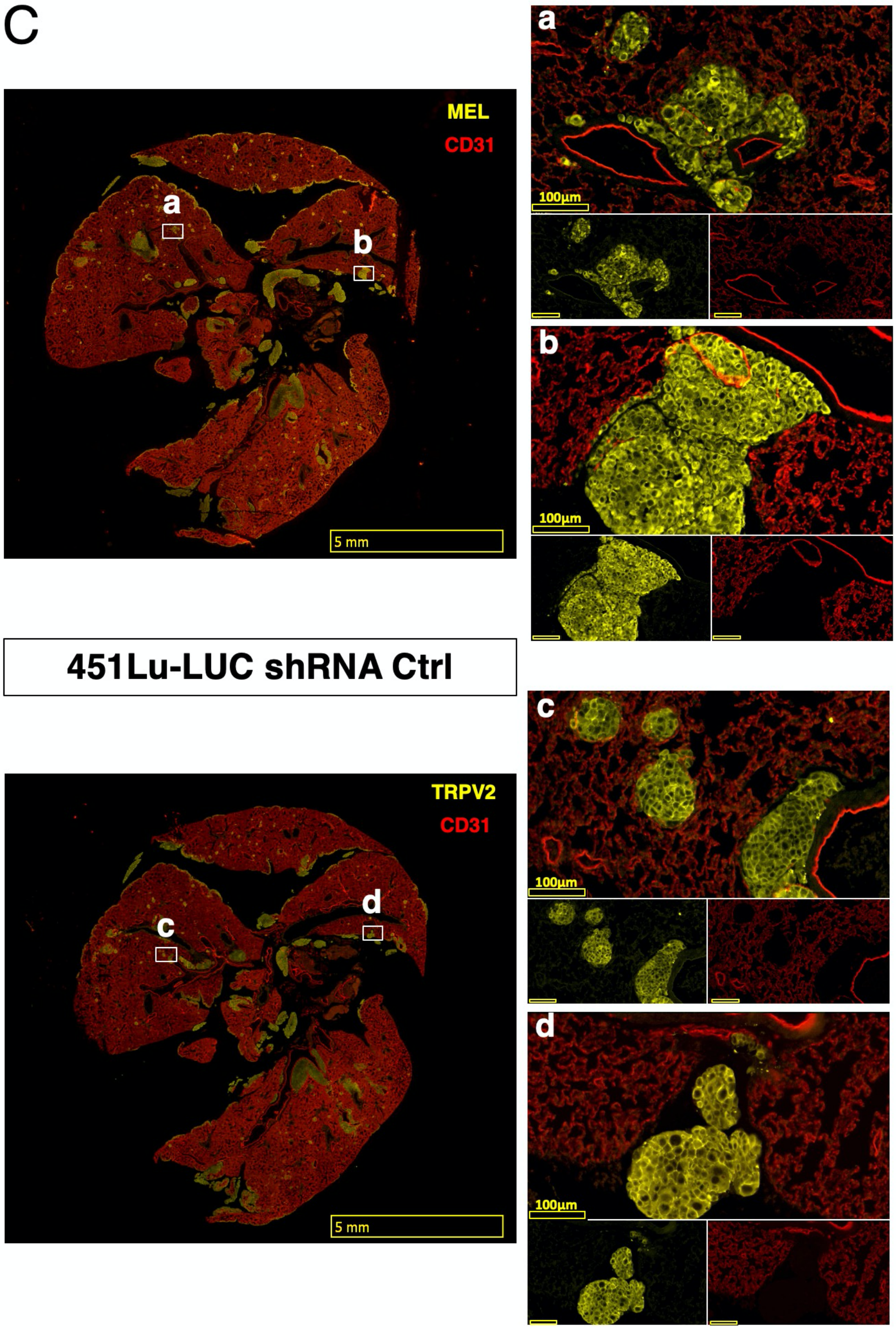
MEL
CD31
a
b
5 mm
451Lu-LUC shRNA Ctrl
TRPV2
CD31
c
d
5 mm
a
100µm
b
100µm
c
100µm
d
100µm

**Supplemental Figure 11. Direct comparison of the *in vivo* metastatic potential of TRPV2-modulated melanoma cells in xenografted zebrafish**

(A) Schematic representation of the experimental strategy used to evaluate the impact of TRPV2 repression in human melanoma cells on micrometastasis formation in xenografted zebrafish. As control- and TRPV2-shRNA expressing WM266.4 cells were both GFP-labeled, control cells were double-stained with the red fluorescent dye CmDil, and equal amounts of both populations were co-injected in the duct of Cuvier of 2 days-old zebrafish embryos. 36h post-transplantation, selected xenografted zebrafish were imaged.

(B) Representative fluorescent microscopy images of zebrafish larvae co-injected with shRNA control (orange) and shRNA TRPV2 (green) melanoma cells. Quantification of tumor burden per embryo shows a significant reduction of the metastatic potential of TRPV2-silenced cells compared to control cells. Eyes autofluorescence was not considered in the quantification. Comparison was done in 9 individual zebrafish larvae (***P<0.005).

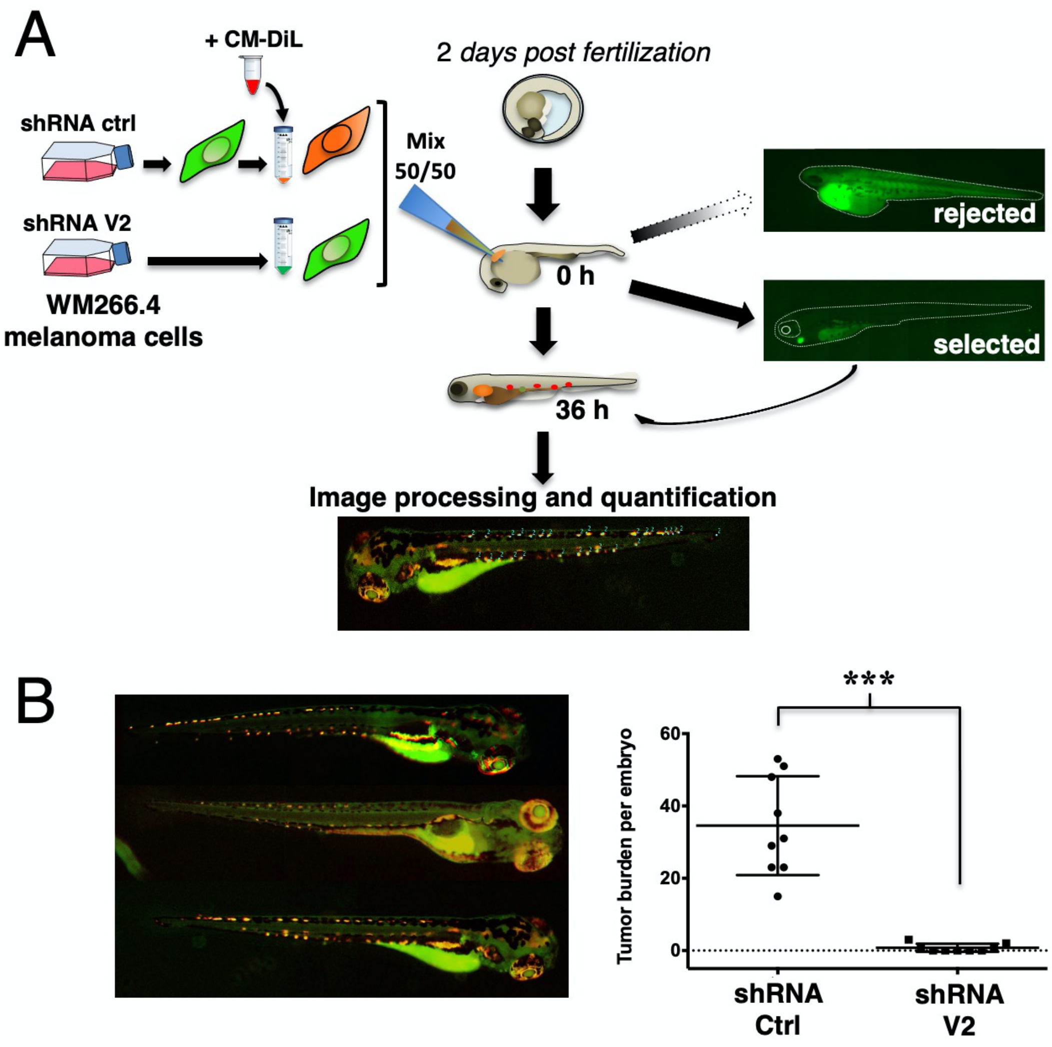

**Supplemental Figure 12 (relative to Figure 7). Validation of the antibodies used to quantify TRPV2 expression in melanoma patient samples**

(A &n B) Specificity of the HPA044993 anti-TRPV2 polyclonal antibody (*Sigma*). Non-transfected HEK cells were used as negative control and human TRPV2 transfected HEK cells as positive control. TRPV2 antibody was tested (A) by western-blot on WM266.4 and 451LU melanoma cells expressing high levels of endogenous TRPV2 or repressed for its expression by specific shRNAs; and (B) by immunohistochemistry on a stage IV melanoma tumor by comparison with its isotype control.

(C) Global picture of the ME1004c melanoma tissue microarray used to analyse TRPV2 expression.

(D) Schematic representation of the image analysis workflow used for TRPV2 staining intensity quantification.

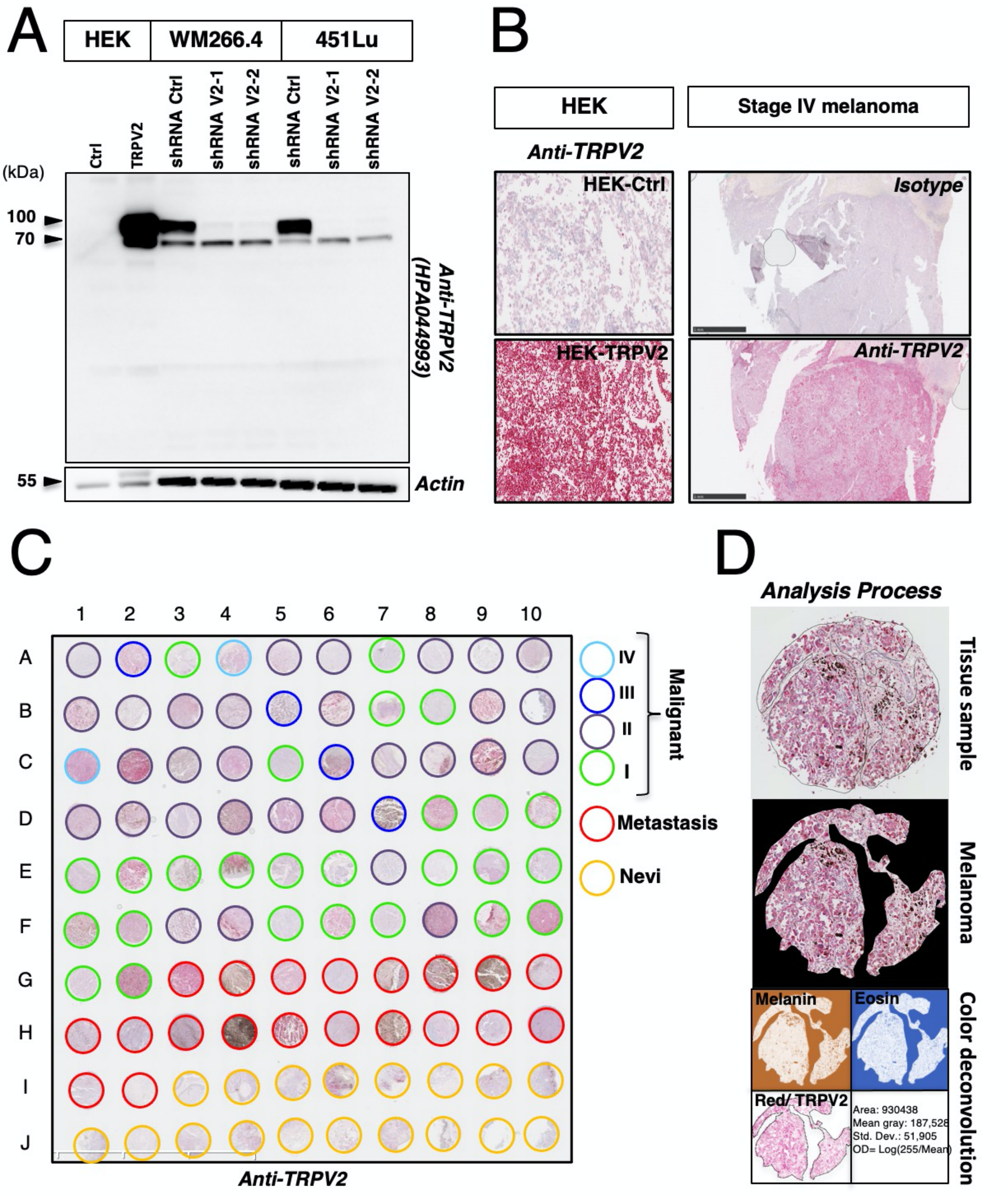

**Supplemental Figure 13 (relative to Figure 7). TRPV2 expression analysis in melanoma patient samples**

(A) RNAseq data-based GEPIA generated dot plot profiling TRPV2 differential expression in the TCGA SKCM tumors cohort (n= 461) *versus* [TCGA normal + GTEx normal] datasets (n= 558). Each dot represents a distinct tumor or normal sample. The method for differential analysis is one-way ANOVA, using disease state (Tumor or Normal) as variable for calculating differential expression: The expression data were first $\log_2$(TPM+1) transformed for differential analysis and the $\log_2$FC was defined as median(Tumor) - median(Normal).

(B) Bar graphs represent TRPV2 protein expression level according to melanoma stages (TNM grading) in the TMA.

(C) Immunohistochemistry staining of TRPV2 on grade I and grade IV melanoma samples issued from Rennes Hospital (CHU) tumor biobank.

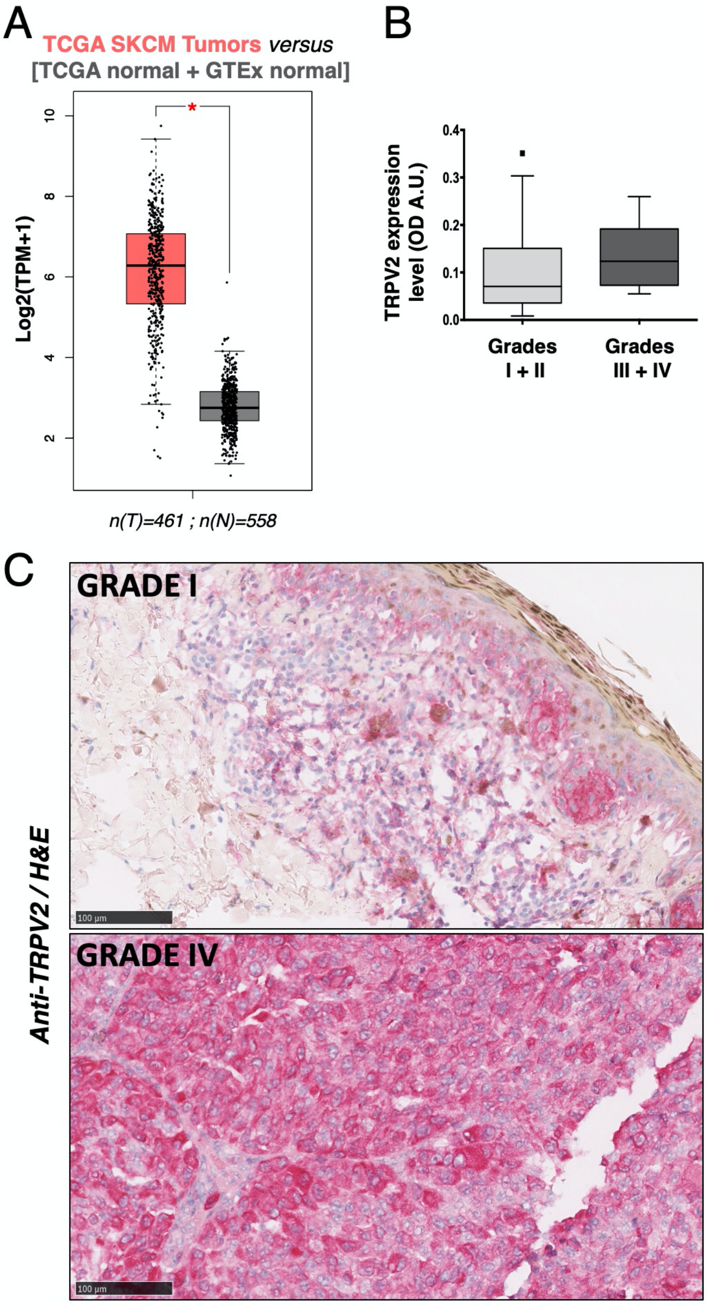

##### Supplemental Movie (relative to Figures 3 and S4)

**Simultaneous live cell imaging of control and TRPV2-silenced melanoma cells.** WM266.4 cells transduced with control (red) or TRPV2 (green) shRNAs placed in an Ibidi migration chamber with a 0-5% FCS gradient and tracked for 12h.
